# Bayesian combination of mechanistic modeling and machine learning (BaM^3^): improving personalized tumor growth predictions

**DOI:** 10.1101/2020.05.06.080242

**Authors:** Pietro Mascheroni, Symeon Savvopoulos, Juan Carlos López Alfonso, Michael Meyer-Hermann, Haralampos Hatzikirou

**Affiliations:** Braunschweig Integrated Centre of Systems Biology and Helmholtz Centre for Infectious Research, 38106 Braunschweig, Germany; KU Leuven, Department of Chemical Engineering, Celestijnenlaan 200F, 3001 Leuven, Belgium; Centre for Individualized Infection Medicine, 30625 Hannover, Germany; Institute for Biochemistry, Biotechnology and Bioinformatics, Technische Universität Braunschweig, 38106 Braunschweig, Germany; Mathematics Department, Khalifa University, P.O. Box 127788, Abu Dhabi, UAE; Centre for Information Services and High Performance Computing, TU Dresden, 01062 Dresden, Germany

## Abstract

In clinical practice, a plethora of medical examinations are conducted to assess the state of a patient’s pathology producing a variety of clinical data. However, exploiting these data faces the following challenges: (C1) we lack the knowledge of the mechanisms involved in regulating these data variables, and (C2) data collection is sparse in time since it relies on patient’s clinical presentation. (C1) implies that only a small subset of the relevant variables can be modeled by virtue of mathematical modeling. This limitation allows models to be effective in analyzing the qualitative dynamics of the system, but limits their predictive accuracy. On the other hand, statistical learning methods are well-suited for quantitative reproduction of data, but they do not provide mechanistic understanding of the investigated problem. Moreover, due to (C2) any algorithm is challenged in learning the corresponding disease dynamics. Herein, we propose a novel method, based on the Bayesian coupling of mathematical modeling and machine learning (BaM^3^), aiming at improving individualized predictions by addressing the aforementioned challenges. As a proof of concept, we evaluate the proposed method on a synthetic dataset for brain tumor growth and analyze its performance in predicting two major clinical outputs, namely tumor burden and infiltration. The BaM^3^ method results in improved predictions in almost all simulated patients, especially for those with a late clinical presentation. In addition, we test the proposed methodology in two settings dealing with real patient cohorts. In both cases, namely cancer growth in chronic lymphocytic leukemia and ovarian cancer, BaM^3^ predictions show excellent agreement with reported clinical data.

## Introduction

Advances in patient care have led to the availability of large amounts of data, generated by typical examinations, such as blood sample analysis, clinical imaging (e.g. CT, MRI) and biopsy sampling, as well as by innovative ‘-omics’ sequencing techniques [1, 2]. Such clinical data are the cornerstone in the practice of personalized medicine and specifically in the field of Oncology [3, 4]. However, this abundance of information comes with multiple issues related to data exploitation and synthesis towards the prediction of pathology dynamics. In particular, we identify the following two major challenges: (C1) First, knowledge of the regulatory mechanisms underlying clinical data is largely lacking, and (C2) patient data collection is usually sparse in time, since patient clinical visits/examinations are a limiting factor.

Regarding the challenge (C1), scientists have been long supported by the use of mathematical modeling as a tool to identify causal relationships in the experimental and clinical data, particularly in cancer treatment [5, 6, 7]. Mathematical models allow to propose and test biological hypotheses, analyze the sensitivity of observables with respect to biological parameters, and provide insights into the mechanistic details governing the phenomenon of interest [8, 9, 10]. Although these models can be extremely powerful both in predicting system responses and suggesting new experimental directions, they require adequate knowledge of the underlying biological mechanisms of the analysed system. Typically, this knowledge is not complete, and only for a limited portion of the involved variables the corresponding mechanistic interactions are sufficiently known. Therefore, even though mathematical models provide a good description of a simplified version of the associated system dynamics, they do not always allow for accurate and quantitative predictions.

On the other hand, machine learning techniques are suitable to deal with the inherent complexity of biomedical problems, but without caring for the knowledge of the underlying interactions [11]. While mathematical models rely on causality, statistical learning methods identify correlations among data [12]. This approach allows to systemically process large amounts of data and infer hidden patterns in biological systems. As a consequence, machine learning-based techniques can provide valuable predictive accuracy upon sufficient training, but do not typically allow for any mechanistic insight into the investigated problem [13]. The overall understanding of the fundamental system dynamics becomes almost impossible, as the chance to generalize the ‘learnt’ system behavior. The latter issue is further exacerbated by the (C2) challenge that has to be faced, related to the sparseness of clinical data. In particular for a single patient, such information is only available at a few time points, corresponding to clinical presentation.

To face the two mentioned challenges, we propose a novel Bayesian method that combines mathematical modeling and statistical learning (BaM^3^) to provide individualized predictions. As a proof of concept, the proposed method is tested on a synthetic dataset of brain tumor growth. We analyze the performance of the new approach in predicting two relevant clinical outcomes, namely tumor burden and infiltration. When comparing predictions from the mechanistic model with those from the BaM^3^ method we obtain improved predictions for the vast majority of virtual patients. We also apply the approach to a clinical dataset of patients suffering from chronic lymphocytic leukemia. The BaM^3^ method shows excellent agreement between the predicted clinical output and the reported data. Finally, as an additional test case, we show how the proposed methodology can be used to assess the time-to-relapse in a dataset of ovarian cancer patients.

## Results

We introduce the key ideas of the proposed methodology in the context of brain tumor growth, leaving the full derivation of the equations and their general form to the Supplementary Information (SI).

Gliomas are aggressive brain tumours generally associated with low survival rates [14]. One of the most important hallmarks of this type of tumors is its invasive behaviour, combined with a marked phenotypic plasticity and infiltrative morphology [15]. The clinical needs led to the development of several mathematical models to support clinicians in the treatment of the disease [16]. As a first test case, we synthetically generate a dataset of glioma patients using a system of Partial Differential Equations (PDEs) recently published [17, 18]. This complex mathematical model (’full model’, in the following) provides a set of *in silico* patients, which represents our synthetic reality and serves as a benchmark to evaluate the performance of the proposed BaM^3^ method. Our goal is to obtain a personalized prediction of the clinical observables of the patients, combining their ‘modelable’ and ‘unmodelable’ information. A simplified mathematical model (with respect to the full model used to generate the patients) is used to generate predictions of clinical outputs starting from the modelable variables. In turn, a machine learning algorithm produces predictions of the same clinical outputs leveraging on the information contained in the unmodelables. As displayed in Fig.1, the core idea of the BaM^3^ method is to use the results of the mathematical model to guide the predictions of machine learning. In more technical terms, the probability distribution function (pdf) obtained from the mathematical model (‘model-derived pdf’, in the following) works as a Bayesian prior that multiplies the pdf obtained from a nonparametric regression algorithm (‘data-derived pdf’). The product of these two pdfs returns an estimate of the pdf for the clinical outputs of interest. More details about the formal definition of the BaM^3^ method and the mathematical details are available in the Methods and SI.

**Figure 1:**
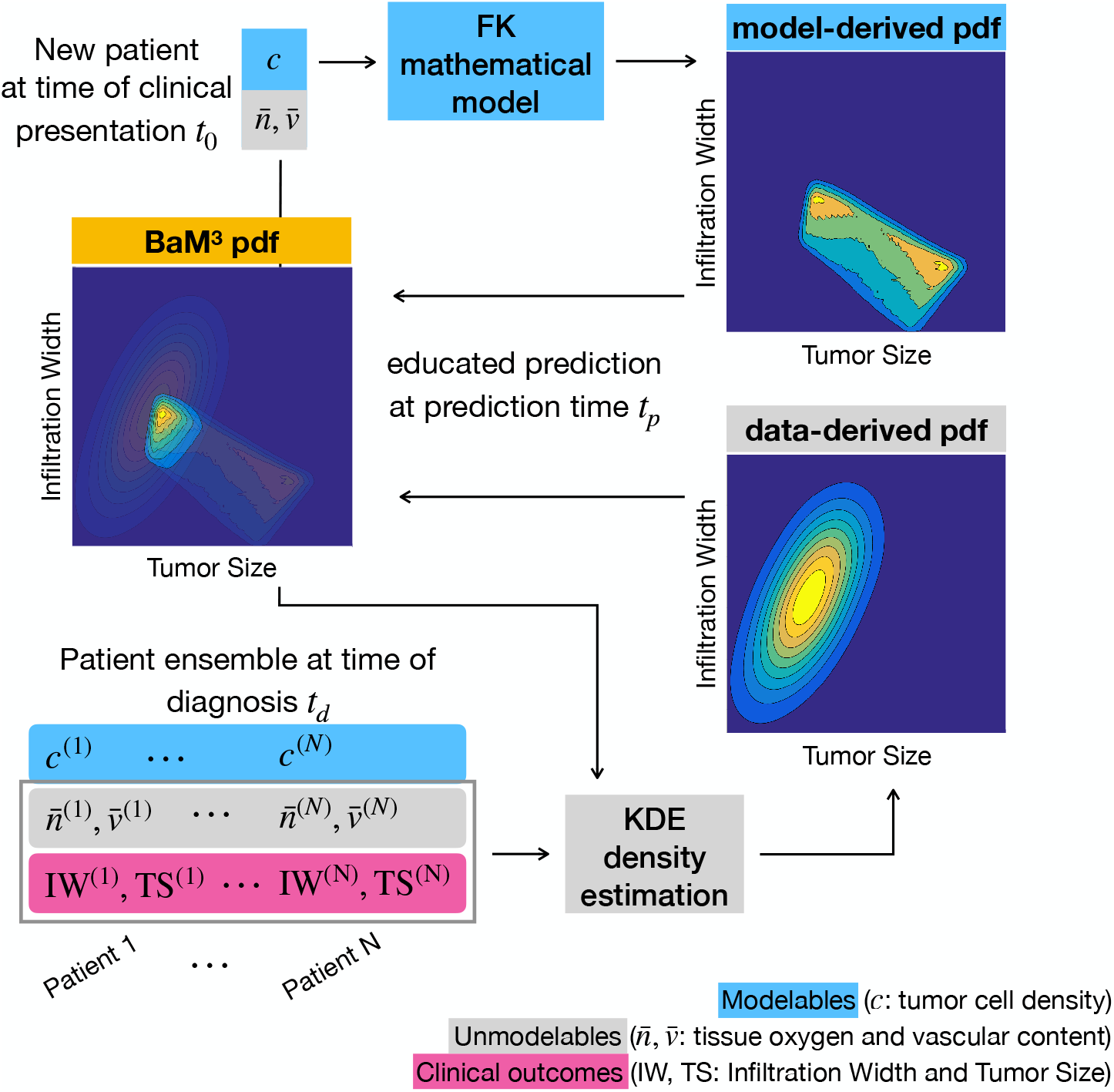
Application of the BaM^3^ methodology to the prediction of glioma growth. Given a new patient characterized by modelable (*c*) and unmodelable 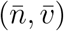 variables at the clinical presentation time *t*_0_, the goal of the BaM^3^ approach is to find an estimate for the clinical observables IW and TS at the prediction time *t_p_*. The modelable variable is used in the FK mathematical model to generate the model-driven pdf. In turn, the unmodelable variables are injected into a density estimation method, which contains the information about the patient ensemble at the diagnosis time *t_d_*. This step provides the data-driven pdf, which is used to correct the predictions from the mathematical model. Application of the BaM^3^ method results in a new pdf for the clinical observables at the prediction time *t_p_*.

### Improving predictions of synthetic glioma growth

For this first test case we deal with two clinical observables, i.e. the Tumor Size (TS) and Infiltration Width (IW). The first quantity is related to tumor burden, whereas the second accounts for tumor infiltration in the host tissue. The modelable variable is the tumor cell density *c*, whereas we consider the amount of oxygen 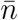 and vasculature 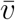 in the tissue to be the patients’ unmodelables - see the Methods. Then, given the patient-specific modelable and unmodelable variables at the clinical presentation time *t*_0_, the BaM^3^ method produces the probability of observing certain values of the clinical outputs at a specific prediction time *t_p_* - see Fig. 1. The data-derived pdf is obtained through a normal kernel density estimator (KDE) [19, 20, 20, 21] incorporating the information about the patient ensemble. The latter is generated from the full model, at the diagnosis time *t_d_*. Then, the model-derived pdf is calculated using the simplified mathematical model for each patient. In particular, we use the Fisher-Kolmogorov (FK) model [22] to produce a map of possible IW and TS starting from the tumor cell density of each virtual patient (see the Methods for further information about the KDE and modeling steps).

Figs. 2**A-C** show the results of applying the BaM^3^ method to a representative patient. We select a clinical presentation time *t*_0_ = 24 months and a time of prediction *t_p_* = 9 months. The model-derived pdf obtained from the FK model is shown in Fig. 2**A**. Interestingly, the prediction of the model in that particular case shows two peaks, one with low TS and high IW and another with opposite properties. We calculate the expected values of TS and IW from the pdf obtained with the FK model and compare it to the ‘true’ values given by the full model. As shown in the plot, for this patient the presence of a bimodal distribution shifts the expected values far from the true ones. We enforce the BaM^3^ method making use of the probability calculated from the KDE, shown in Fig. 2**B**. The latter pdf takes into account the correlations between the clinical outputs and the unmodelable variables present in the patient ensemble. For this patient, the unmodelable distribution selects the probability mode closer to the true IW and TS values, as displayed in Fig. 2**C** (another example for a different patient is given in Fig. S3 in the SI). These results evidence the ability of the proposed method to correct the predictions obtained by using exclusively the mathematical model, and to produce an expected value of the pdf that is closer to the ground truth.

**Figure 2:**
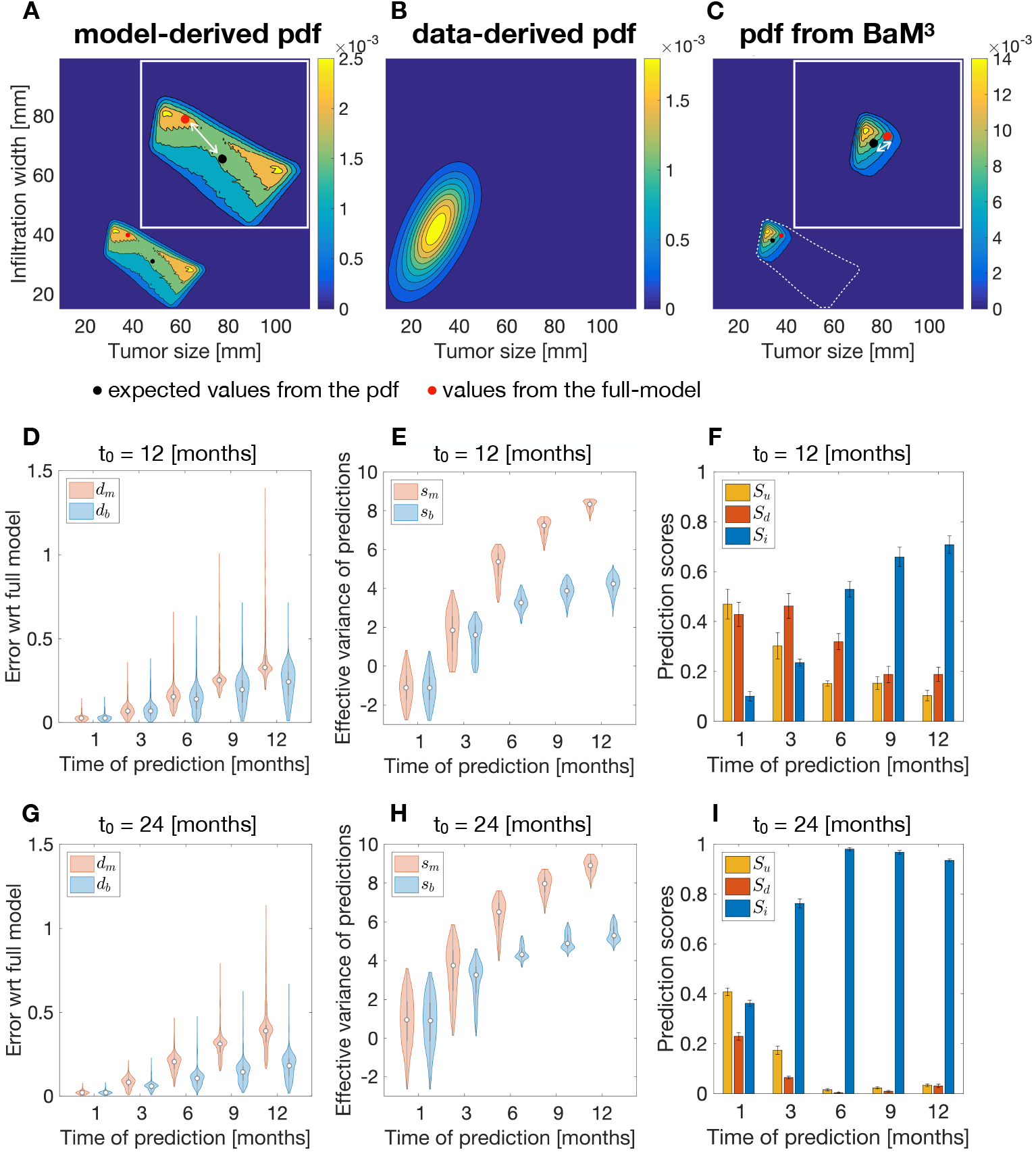
**A-C** Application of the BaM^3^ method to a representative patient. The clinical presentation time *t*_0_ is of 24 months and the prediction time *t_p_* is 9 months. **A** Pdf obtained from the FK model, plotted over the (TS, IW) space. The inset shows an enlargement of the probability region, in which the white arrow marks the error between the clinical output predicted by the full model (red dot) and the expected value of the distribution (black dot). **B** Data-derived pdf for the specific patient calculated through the kernel density estimator trained over the patient ensemble. **C** Probability distribution obtained from the BaM^3^ method. The dashed line shows the original probability obtained from the FK model. Relative errors between the clinical outputs obtained from the full model and the expected values given by the FK model (*d_m_*), and after the application of the BaM^3^ method (*d_b_*), for a clinical presentation time of 12 and 24 months (**D**,**G**, respectively). Effective variance of the predictions for the corresponding clinical presentation times (**E**,**H**, respectively). Prediction score, as the ratio of cases for which the BaM^3^ method improved (*S_i_*), did not change (*S_u_*) or deteriorated (*S_d_*) the model predictions. The error bars show the variation of the results by replicating the method over 10 different sets of N=500 patients.

We apply the BaM^3^ method for two clinical presentation times at 12 and 24 months, and compare its outcomes with those provided by the full model at increasing prediction times (Fig. 2**D,I**). For each patient we calculate the relative error between the predicted clinical outputs obtained from the full model and the expected values of the pdf calculated from the FK model (*d_m_*) and after implementing the BaM^3^ method (*d_b_*). These nondimensional errors are calculated as

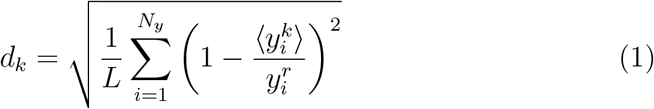

where 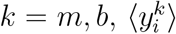 are the expected observable values (*i* =IW, TS) calculated from the FK model and the BaM^3^ method, and 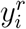 are the observable values obtained from the full model. We calculate the errors *d_m_* and *d_b_* for each patient at different prediction times. Then, we compare the corresponding errors and evaluate if the BaM^3^ method improved, deteriorated or left unchanged the prediction from the FK model, i.e. *d_b_ < d_m_*, *d_b_ > d_m_* or *d_b_ ~ d_m_*, respectively (see Methods). We denote the ratio of improved, unchanged and deteriorated cases with respect to the total number of simulated patients as *S_i_*, *S_u_* and *S_d_*, respectively.

Both the relative errors *d_m_* and *d_b_* increase for increasing prediction times, as shown in Fig. 2**D,G** for the two clinical presentation times considered. However, after applying the BaM^3^ method the errors decrease, especially at later times. In general, it is possible to notice an improvement both in terms of median values and sparseness of the data. Interestingly, the relative error obtained from the BaM^3^ method increases at a lower rate if compared to the relative error obtained from the FK model.

We also calculate the effective variance of the predictions as the logarithm of the determinant of the covariance matrices relative to the model and BaM^3^ pdfs, (identified by *s_m_*, *s_b_* respectively - see Methods). This quantity reflects the spreading of the pdfs over the (TS, IW) plane, with higher values denoting more uncertainty in the predictions. For both clinical presentation times (Fig. 2**E,H**) the BaM^3^ method provides thinner pdfs, more centered around their expected value with respect to the FK model-derived case.

Finally, the stacked bars in Fig. 2**F,I** show that BaM^3^ performs well at later prediction times and remarkably well (improvement ratio *S_i_* close to 1), especially at the latest clinical presentation time *t*_0_ = 24 months. For *t*_0_ = 12 months (Fig. 2**F**) the proposed method is not able to improve predictions until a prediction time of 6 months. Then, for *t_p_* = 6, 9 and 12 month the advantages of using BaM^3^ over the FK model are unambiguous. On the other hand, for the clinical presentation time of 24 months (Fig. 2**I**) both *S_u_* and *S_d_* decrease significantly for prediction times equal or greater than 3 months. The ratio of improved cases *S_i_* reaches almost 100% at each of the last three prediction times, clearly overcoming the results of the FK model. The error bars in Fig. 2**F,I** denote the variability in the results that is obtained by replicating the study 10 times, each with *N* = 500 randomly generated patients. Fig. S4 in the SI shows similar results when decreasing the number of patients. Notably, the scores for *N* = 500, 250, 100 and 50 are very close, slightly improving with increasing the number of patients. The variability in the 10 replicates also decreases for higher values of *N*.

We also calculate the prediction scores using the distribution mode to generate the scores instead of the expected value (see Fig. S5). When the pdfs display multiple maxima we consider the average of the relative errors between the values of the full model (i.e. the synthetic reality) and the different peaks. The performance of the BaM^3^ method sensibly degrades with respect to using the expected value. Improvement in predictions is observed only for later clinical presentation and prediction times.

In summary, the BaM^3^ method is able to correct the FK model predictions for most of the patients, particularly at later clinical presentation and prediction times. The improvement in the prediction occurs by (i) decreasing the median relative error between expected observable values and ground truth, (ii) decreasing the rate at which the error increases with prediction time, anddecreasing the variance associated with the probability distributions.

#### BaM^3^ performance depends on the clinical output

Even though the BaM^3^ method performs well for the majority of patients, there are some cases for which it fails to improve the predictions of the mathematical model. We analyze the failure cases by splitting the errors *d_m_* and *d_b_* into the two partial errors

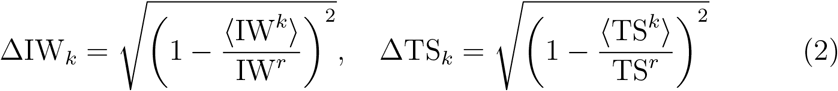

where *k* = *m, b*, 〈IW^*k*^〉 and 〈TS^*k*^〉 are the expected values of the clinical outputs obtained from the mathematical model and BaM^3^ pdfs (*k* = *m, b*, respectively), and IW^*r*^, TS^*r*^ are the values of these quantities from the full model. Fig. 3 shows how these partial errors are distributed over the presentation and prediction times. The dashed line in the plots highlights the neutral boundary, where the partial errors of the FK model and BaM^3^ method are equal. Above this line, the proposed BaM^3^ method deteriorates the model predictions, whereas under that line the BaM^3^ method improves predictions. The red dots in the scatter plots represent the patients for which the BaM^3^ method fails (‘failure cases’, in the following). After a prediction time of one month, in which a characteristic pattern is not evident, the plots highlight that failure cases are generally associated with regions where the BaM^3^ method under-performs to the FK model with respect to TS (ΔTS_*b*_ > ΔTS_*m*_). Interestingly, the same failure cases belong to regions in which ΔIW_*b*_ < ΔIW_*m*_: the BaM^3^ method is improving the IW predictions and at the same time deteriorating the TS predictions. This happens for both *t*_0_ = 12 and 24 months, however the number of failure cases is considerably higher for the earlier presentation time. For the specific case under consideration, lower performance of the BaM^3^ method is therefore associated to its inability in correcting the FK model predictions for TS, with a tendency that improves for the later presentation time due to strong corrections for IW.

**Figure 3:**
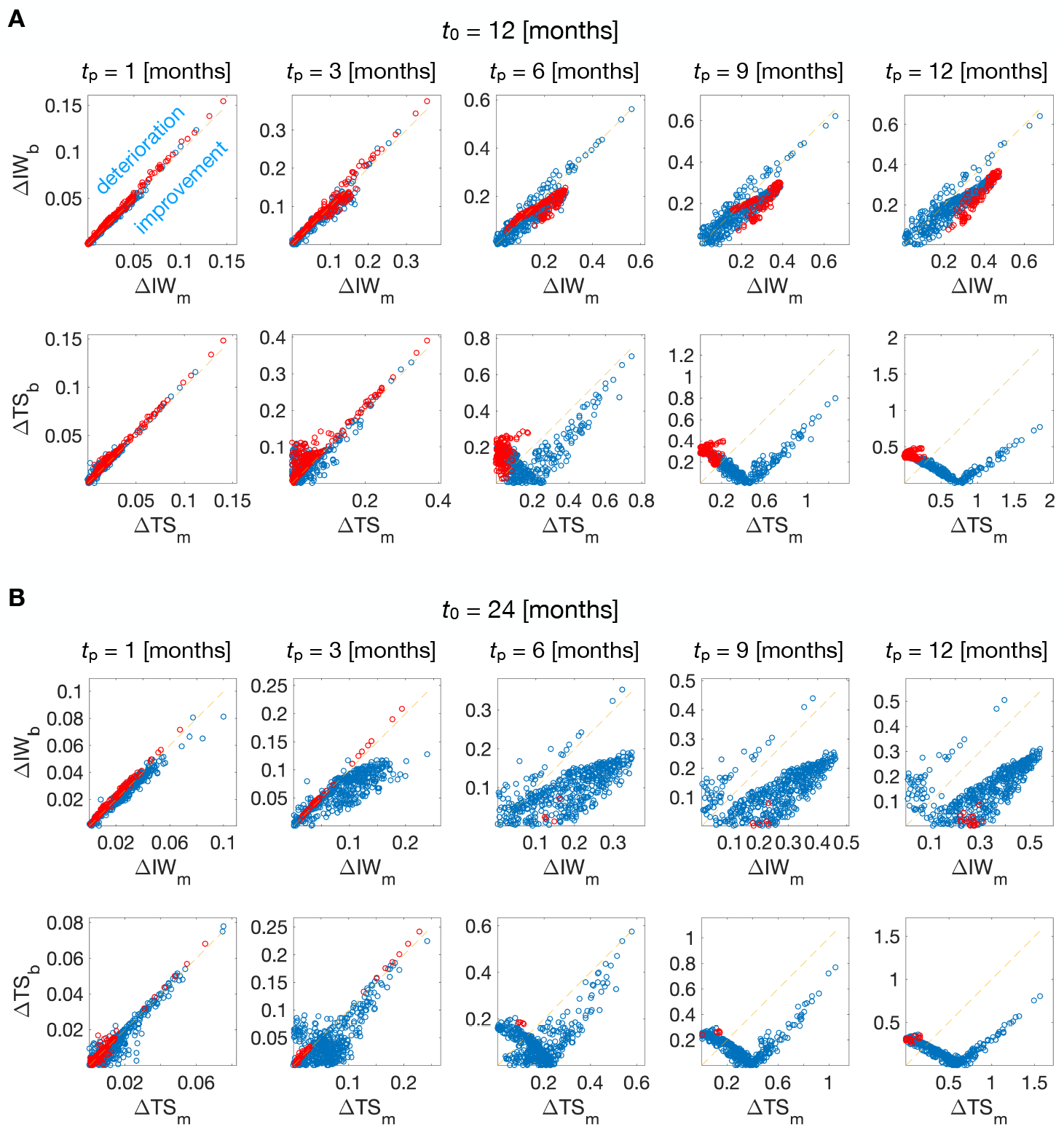
Scatter plots of the partial errors ΔIW_*i*_, ΔTS_*i*_ over different prediction times *t_p_* (*i* = *m, b*, for the quantity related to the mechanistic model or the BaM^3^ method, respectively). Results for presentation times of *t*_0_ = 12 months (**A**) and *t*_0_ = 24 months (**B**) are shown. Each red dot corresponds to a patient for which the BaM^3^ method fails in improving the FK model predictions. Regions over the dashed line correspond to areas in which the BaM^3^ method deteriorates the FK model predictions, while regions under that line show areas in which it improves the FK predictions.

#### Transient behavior of the unmodelable distribution is associated to limited improvements

To investigate the reasons for the poor performance of the BaM^3^ method in improving the predictions for one of the clinical observables, we analyze the behavior of the pdf arising from density estimation, i.e. the data-derived pdf.

Fig. 4 shows the temporal evolution of this quantity for different clinical presentation times *t*_0_. Fig. 4**A** shows a plot of the unmodelable pdf for a representative patient over the clinical output space. From a pdf that covers a limited region in the (IW,TS) plane, the probability distribution spreads over a broader area as the presentation time increases. The center of mass of the distribution, however, tends to converge to a more specific region as time progresses. This is more evident in Fig. 4**B,C**, showing the marginal probabilities for IW and TS, calculated from the distribution in Fig. 4**A**. The marginal distributions become broader for both IW and TS, but in the first case their peak stabilizes at later *t*_0_ times. On the contrary, the peak of the marginal probability for TS moves towards larger values at higher times. To quantify this behavior across the different patients we then evaluated the degree of overlap between the marginal probabilities at two subsequent *t*_0_. Results from this calculation are plotted in Fig. 4**D,E** for the overlap between the distributions at presentation times *t*_0_ of 12 and 18 months and between 24 and 30 months. Here, the degree of overlap is calculated as the area of overlap for the IW and TS marginal distributions. Values close to one represent maximum overlap, whereas values near zero are associated to poor overlap between the two marginal pdfs. In a rough approximation, when this overlap score is high the marginal pdf is close to a steady state (since the pdf has not moved over time), and viceversa. For the earlier times in Fig. 4**D** the patients are mostly scattered along a line of increasing IW_*r*_ and TS_*r*_ with points where the overlap is poor (close to 0.4 in certain regions). On the other hand, for the later case in Fig. 4**E** the patients are shifted towards higher values of overlap. Moreover, a horizontal line of high overlap for the IW output is visible for a large patient ensemble, pointing to a stabilization towards a steady state for the IW at later presentation times. This explains the lower BaM^3^ method performances at *t*_0_ = 12 months, since the pdf from the KDE that should correct the model predictions is projecting the model pdf over (IW,TS) values that are outdated, far from the steady state. The situation improves for the case of *t*_0_ = 24 months: even though the correction of the BaM^3^ method for the TS might be wrong is some cases, the pdf for the IW has stabilized and points towards the correct value. In most of the cases the correction for the IW outperforms the one for the TS which leads to a general improvement of predictions by the BaM^3^ method.

**Figure 4:**
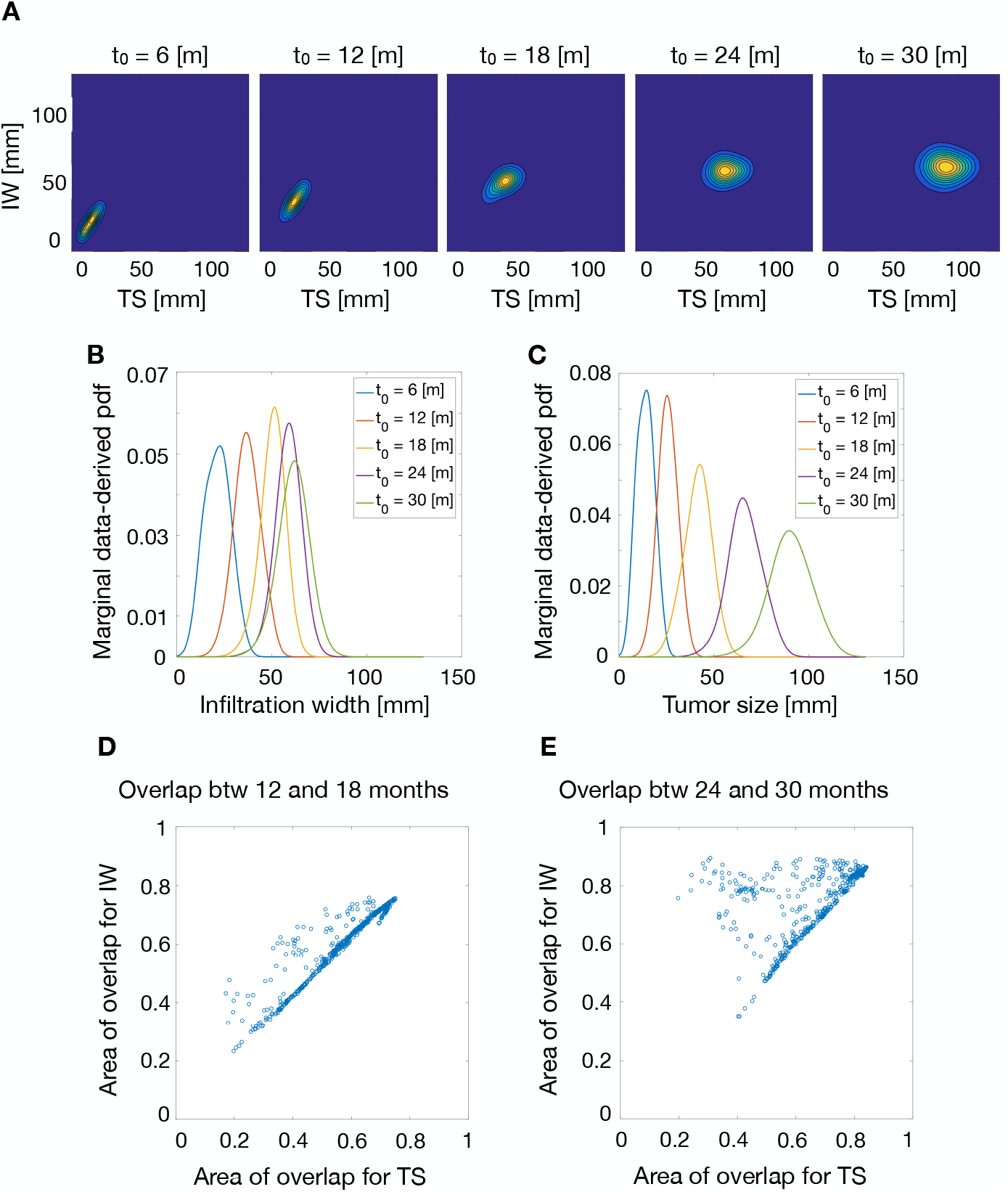
Temporal evolution of the unmodelable pdf from the KDE step. **A** Data-derived pdf for a representative patient at different clinical presentation times, plotted over the (IW,TS) space. Marginal probabilities from the plots in **A**, showing the evolution of the probability distribution for IW (**B**) and TS (**C**). Scatter plots representing the degree of overlap between the marginal probabilities for IW and TS between clinical presentation times *t*_0_ of 12 and 18 months (**D**) and of 24 and 30 months (**E**).

#### Outlier patients challenge the method’s performance

To explain the different behavior for IW and TS, we investigate the distribution of the failure cases over the full model parameter space. In general, the BaM^3^ method performs poorly for those patients that are at the extremes of the parameter space, who represent outlier patients. When plotting the patients in a scatter plot over cell motility and proliferation rate (Fig. 5), the points with high motility - high proliferation rates and high motility - low proliferation rates witness the highest number of failure cases for both clinical presentation times *t*_0_ of 12 and 24 months. We checked for the distribution of failure cases also for the other model parameters, but no particular pattern was evident (see Fig. S6). Notably, patients falling into these high motility - high/low proliferation regions show the highest values for IW and TS (see [17] and Fig. S7). Highly invasive and massive neoplasms are inadequately described by the pdf from the KDE, as they represent the extreme cases of the probability distribution. As a result, the FK model performs better in predicting the clinical outcomes with respect to the BaM^3^ method, since in the latter the correction from the dataset points towards smaller values of IW and, especially, TS.

**Figure 5:**
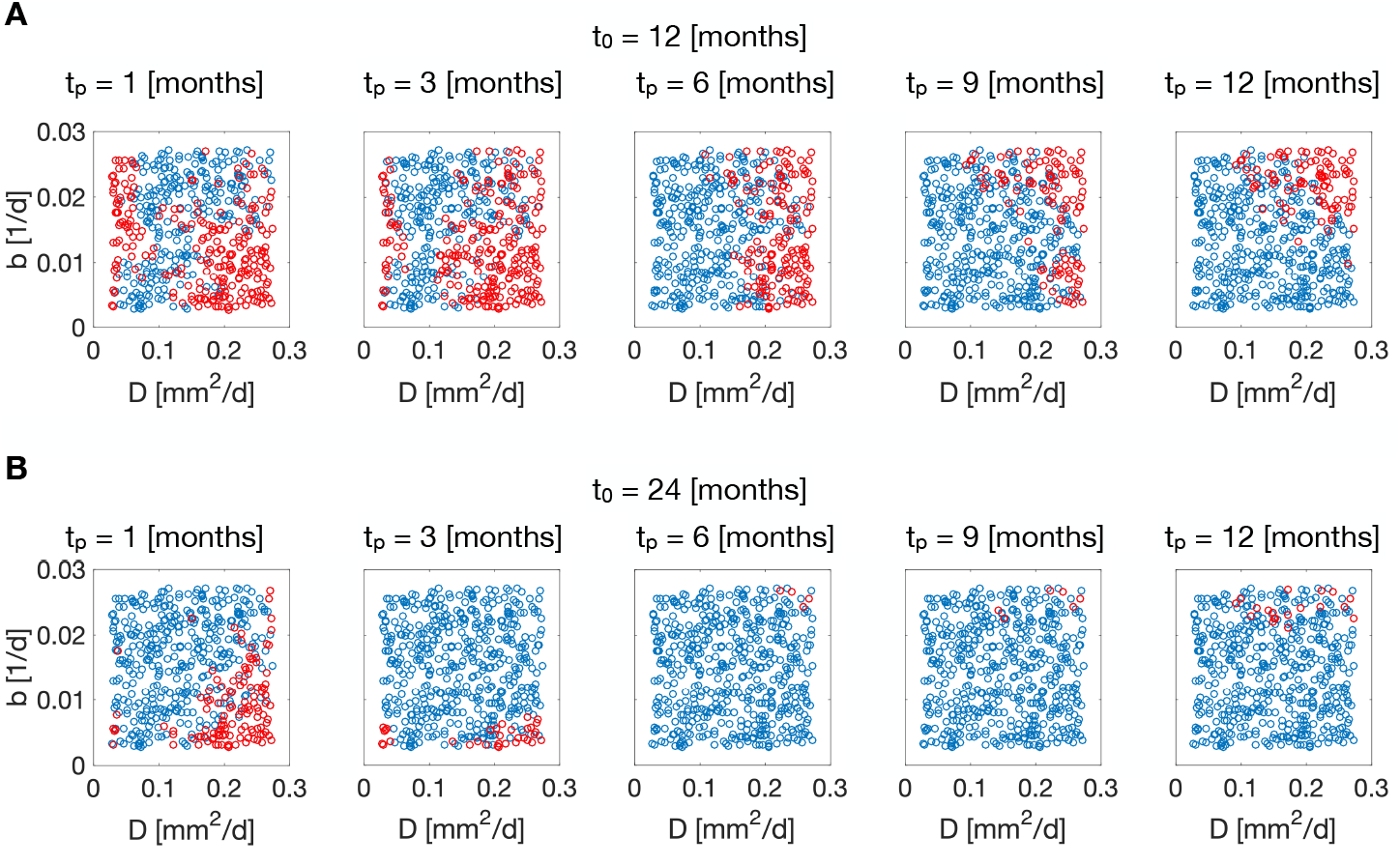
Scatter plots showing the distribution of failure cases (red dots) over the parameter space at different presentation and prediction times. *D*: tumor cell motility; *b*: tumor cell proliferation rate; *t*_0_: clinical presentation time; *t_p_*: prediction time.

### Applying the method to real CLL patients: the effect of unmodellables

In addition to the proof of concept applied to *in silico* data, we test the BaM^3^ methodology on a cohort of real patients suffering from Chronic Lymphocytic Leukemia (CLL). This cancer involves B cells and is characterized by the accumulation of lymphocytes in the blood, bone marrow and secondary lymphoid tissues [23]. In the past, CLL was considered to be a homogeneous disease of minimally self-renewing B cells, which accumulate due to a faulty apoptotic mechanism. This view was questioned by recent findings, suggesting a more heterogeneous neoplastic population continuously interacting with its microenvironment [24, 25, 26, 27]. Accumulation of leukemic cells occurs because of survival signals originating from the external environment and interacting with leukemic cells through a variety of receptors. The nature of this cross-talk with the environment is a current matter of research, featuring *in vitro* as well as *in vivo* experiments. One of the most significant experiments involving human patients was that of Messmer et al [28]. Messmer and his co-workers inferred the kinetics of CLL-B cells from a group of patients through non-invasive labeling and mathematical modeling. Their investigation was quite thorough and involved the collection of several quantities related to patients’ personal data (gender, age, etc.) and status of the disease (years since diagnosis, treatments, mutation status, etc.). They measured the fraction of neoplastic labeled cells in the blood of the patients, and fitted an Ordinary Differential Equation (ODE) compartmental model to the dynamics that they observed. The model included three parameters, i.e. the daily water exchange rate (*f_w_*), the B cell birth rate (*b*), and the relative size of the blood compartment (*v_r_*).

We use the same model as Messmer and colleagues as the input for the BaM^3^ method, but discard the patient-specific fitting provided in their publication. Our aim is to show that, even when an individualized model parametrization is unknown, coupling the information given by the unmodelables can provide good patient specific predictions. To accomplish this, we run simulations over uniform parameter ranges to obtain the pdf of the labeled cell fraction at day 50 (*f*_50_), which is also the sole modelable variable in this dataset (see the Methods and Fig. S8). Then, we incrementally select one to four unmodelable variables from the patients’ dataset and build the data-derived pdf using the same KDE method as in the previous *in silico* example. The BaM^3^ method couples the two prediction distributions to obtain the pdf for the clinically relevant output (see Fig. S9). We show the results of this procedure in Fig. 6**A**, where we compare the BaM^3^ predicted values against the patient *f*_50_ values reported in [28]. The fraction of labeled cells predicted by the BaM^3^ method agrees well with the reported data, especially when we increase the number of unmodelables used for density estimation. The inset shows how the Mean Squared Error (MSE) of BaM^3^ predictions decreases after considering all the possible combinations of unmodelables. Fig. 6**B** shows how the probability distribution generated from the KDE changes for a representative patient. As the number of unmodelables increases, the mode of the distribution shifts towards the correct value of *f*_50_, here denoted by a red dashed line. From Fig. 6**A** it is also possible to note that, even if the majority of points lies close to the perfect prediction line, the predictions of a few patients are significantly mismatched with respect to the corresponding real values. This occurs because these patients belong to the extremes of the parametric space (see Fig. S10). Patients characterized by outliers in their parametriziation are under-represented in the modelable pdf due to the uniform sampling of the parameter space, and it is challenging for the data-derived correction to improve predictions for them.

**Figure 6:**
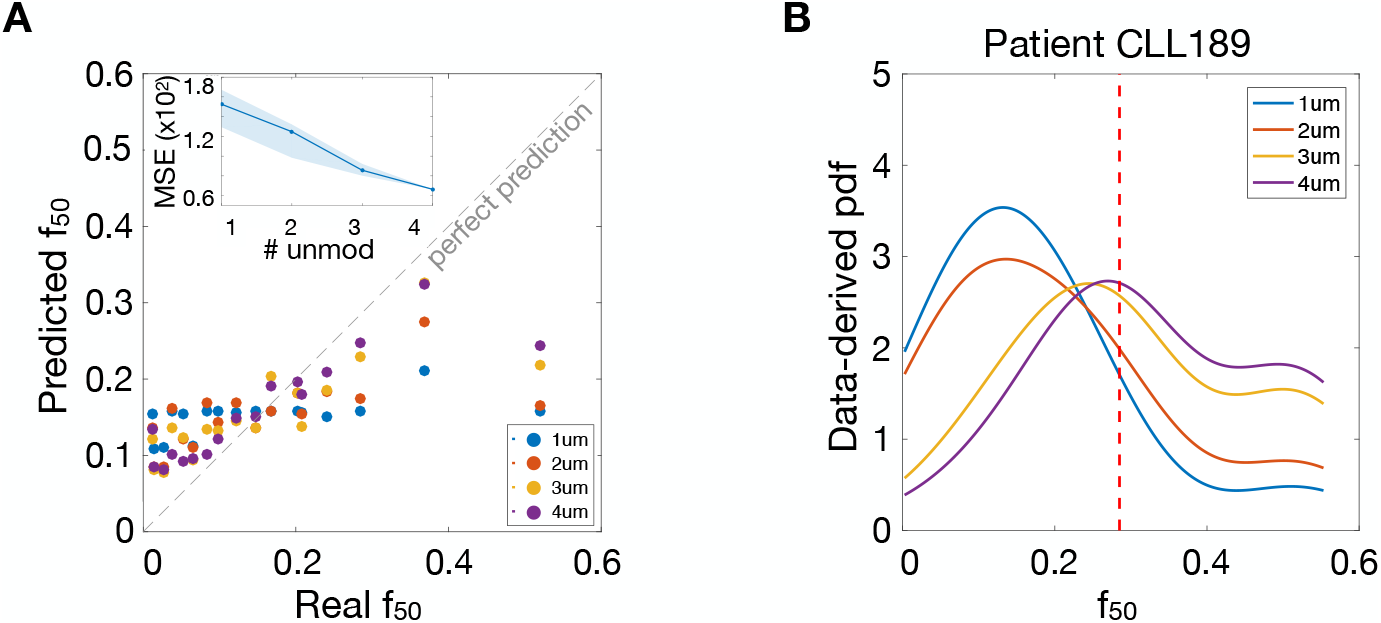
**A** Predicted vs measured values of the fraction of labeled cells at day 50. The colors denote the use of an increasing number of unmodelables in the BaM^3^ method. The inset shows the decrease in Mean Squared Error (MSE) for the BaM^3^ predictions when we consider all the possible combinations of unmodelables. The solid line in the inset refers to the unmodelables used for the scatter plot. Here we included, in order, the CD38 expression, age, growth rate of white blood cells, and V_H_ mutation status as unmodelables. **B** Variation of the data-derived pdf obtained from density estimation when an increasing number of unmodelables is considered for a specific patient. The dashed red line highlights the measured value of *f*_50_ in this patient.

### Prediction of the time-to-relapse on a real ovarian cancer patient cohort: the importance of adequate model parametrization

To provide another application of the BaM^3^ method to a real scenario, we consider the case of patient response to therapy in high-grade serous ovarian cancer (HGSOC). This type of cancer is the most common epithelial ovarian cancer subtype, accounting for 70% to 80% of ovarian cancer–related deaths [29, 30]. In addition, due to treatment resistance, the 5-year survival rate in HGSOC is less than 50% [31, 32]. Indeed, the contribution of resistance mechanisms to tumor relapse after therapy is currently an active matter of research, recently backed up by evolutionary studies [33, 34, 35].

We start from the clinical dataset provided in a recent publication [36], and elaborate a strategy to predict the time-to-relapse (TtR) in ovarian cancer patients which makes use of the BaM^3^ methodology. The database of patients consists of 20 individuals, which are subject to the following treatment schedule (see Fig. S11 in the Supplementary). First, the patients receive neoadjuvant chemotherapy (NACT), consisting of different cycles of carboplatin and paclitaxel chemotherapy. Then, a surgery is performed, followed by other cycles of adjuvant chemotherapy. We propose a low-dimensional mathematical model to predict tumor TtR after treatment for each patient, which takes into account the presence of two cell subtypes. In particular, we include cells that are sensitive or resistant to chemotherapy. In addition, we consider the age of the patient at diagnosis as the unmodelable quantity used by the density estimator. Full details of the model and methodology are available in the Methods and SI.

As in the previous sections, the pdf from the mathematical model is obtained by simulating the latter over the parameter space. In this case, we focus on two parameters, namely the initial fraction of sensitive cells *x*_0_ and the death rate induced by chemotherapy *δ*_0_. First, we consider a uniform distribution of both parameters. We assume *x*_0_ between 0.4 and 0.9, in agreement with the degree of variability reported on the publication from which we take the dataset [36]. Since we lack any information about *δ*_0_, we select a wide range, from 0.1 and 10 d^−1^. This result in an almost uniform pdf from the mathematical model, as shown in Fig. S12. In this condition, the pdf from the model enters the BaM^3^ as an uninformative prior in the Bayesian framework, leaving predictions to rely only on the pdf generated from the density estimation of the unmodellables. Note that, in these settings, BaM^3^ reduces to nonparameteric regression [20]. We calculate the mean squared error (MSE) using the mode of the distributions in the uninformative case - denoted as MSE_un_ - and find MSE_un_ = 38.901 months^2^. As a next step, we use the additional information provided in the dataset to improve the parametrization of the mathematical model. Indeed, the dataset reports the tumor volume before and after the first cycle of therapy, as measured from clinical imaging [36]. We fit the value of *δ*_0_ for each patient and take the mean of all these values as the center of another uniform distribution (the details are in the Methods). We apply the BaM^3^ method using the newly generated pdf from the model and obtain a lower MSE, i.e. MSE_fit_ = 30.895 months^2^ (see Fig. S13). Better parametrization results therefore in improved performace of the method. In addition, by applying BaM^3^ to a better parametrized model allows to obtain improved predictions, as shown in Figure 7. The scatter plot in Figure 7**A** shows reduced errors in BaM^3^ predictions with respect to the ones from the mathematical model or density estimation alone. Also, 7**B** displays the outcome of the method for two representative patients. In both cases, the pdf arising from BaM^3^ has its mode closer to the real TtR (dashed line), with respect to the modes obtained from the model or density estimation pdfs. This shows the potential of the BaM^3^ method, which is able to perform better than the single techniques upon which it is based.

**Figure 7:**
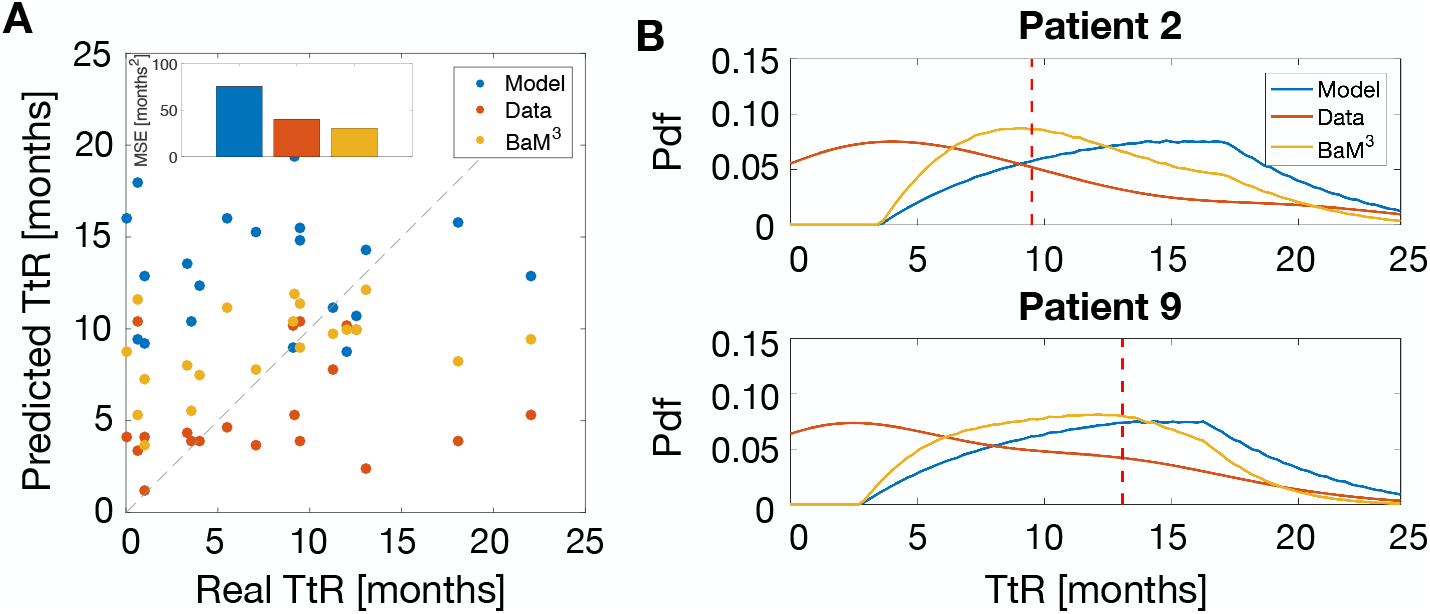
**A** Predicted vs real TtR calculated from the mode of the probability distribution obtained from the mathematical model (Model), density estimation (Data) and the BaM^3^ method. As shown in the inset, the MSE are of 75.674, 40.447 and 30.895 months^2^, respectively. **B** Probability distributions obtained for two reference patients. The red dashed line denotes the real value of TtR for the specific patient. TtR: time-to-relapse of the tumor after therapy.

## Discussion

In the last few years, mathematical modeling and machine learning have emerged as promising methodologies in the biomedical field [37, 38, 39]. However, several challenges persist and limit the prediction accuracy of both approaches. Among these issues, we identified the lack of knowledge of the mechanisms that govern the system under study (C1), and the paucity of time points at which patient information is available (C2), to significantly limit the performance of both mathematical models and machine learning techniques. In this work, we presented a novel method (BaM^3^) to couple mathematical modeling and density estimation in a Bayesian framework. The goal of BaM^3^ is to improve personalized tumor burden prediction in a clinical setting. This coupling allows to address the aforementioned (C1) and (C2) challenges, by exploiting the strengths of the respective methodologies and integrating them in a complementary path.

In particular, our proposed method aspires to solve a dire problem in personalized medicine that is related to the limited time-points of patient data collection. This implies that data assimilation methods, such as Kalman filters or particle filters [40, 41], that require multiple data time-points integrated to a mechanistic model, cannot be generally used. To this regard, the BaM^3^ method can be regarded as an one-step data assimilation method. Compared to other methodologies that combine outputs from mathematical models and measured data - such as Bayesian Melding [42], History Matching [43], Bayesian Model Calibration [44] or Approximate Bayesian Computation [45] - the BaM^3^ method is not interested in parameter estimation to better calibrate the mathematical model. Instead, its goal is to improve predictions of mathematical models empowering them with knowledge from variables that are not usually considered (the ‘unmodelables’, in our framework). This is done without the exact knowledge of the parameters of the mathematical model - indeed we calculated the pdfs of the modelable variables using a uniform sampling of the parameter space. Better estimation of the model parameters improves the outcomes of the method (as shown in the ovarian cancer test case), but it is not required for the methodology to be applied.

First, we tested the BaM^3^ method on a synthetic dataset of patients focusing on tumor growth dynamics. Our approach was able to improve the predictions of a FK model for the majority of the virtual patients, with significant improvements at later clinical presentation times. In addition, we tested the proposed methodology on two clinical datasets related to cancer, concerning tumor growth in leukemia and ovarian cancer patients. We compared the outcomes of the BaM^3^ method to the reported data and found excellent predictive capability. When analyzing the cases for which the performance of the BaM^3^ method was not optimal, we came across some limitations that should be addressed when applying the methodology to real cases.

The first limitation regards the selection of the proper unmodelable variables. These are quantities that cannot be easily mathematically modeled, but can be correlated to the patient clinical outputs. For our proof of concept we selected only a few unmodelables, but in principle multiple quantities could be considered at the same time. Moreover, the most important unmodelables could be selected in a process of feature selection similar to the ones usually adopted in machine learning, providing better accuracy for the predictions [46, 47]. We note as well that the method is open to progress in knowledge: Should an unmodelable variable become modelable because of an increased understanding of the biological mechanisms, this variable can change side and become modelable.

One should also propose an adequate mathematical model that describes the dominant dynamics of the disease, as shown in the last case for ovarian cancer. A better parametrization of the model facilitates the work of density estimation, considerably reducing prediction errors. Not only better model parametrizations are advocated, but also mathematical models that encompass a suitable amount of mechanisms about the phenomenon that is modeled. In the case of ovarian cancer, we show in the SI that a simplified model (with respect to the two cell population presented in the Methods) is not able to provide good predictions when used in the context of the BaM^3^ method (see also Fig. S14).

Care must be taken with the selection of the metric that should be improved by the BaM^3^ method. For the *in silico* case, for example, considering the expected value of the final pdf resulted in better method performance when compared to selecting the pdf mode (see Fig. 2 and Fig. S5). This was probably due to the very similar natures of the FK and full models. Indeed, for lower clinical presentation times the FK model is already ‘primed’ towards the correct solution (in terms of outcomes of the full model); applying the BaM^3^ method might result in adding noise to the FK prediction, degrading the final prediction. However, for some patients the FK model provides pdfs with multiple local maxima, sometimes far away from the full model values. In these instances (see Fig. S3) the BaM^3^ method is able to correct for the correct mode, shifting the pdf to the correct values. Therefore, a good practice would be to try multiple pdf metrics and test the BaM^3^ method on each of them. This would result in a more thorough understanding of the problem, eventually allowing for better predictions.

Another important issue is that the correlations between unmodelables and clinical outputs should be persistent over time, evolving on a timescale that is faster than the dynamics of the problem. In our synthetic dataset this was partially accomplished at later clinical prediction times, especially for the case of the tumor infiltration width. Indeed, the unmodelables variables need to provide as much as possible time-invariant information on the clinical output variables, implying an equilibrated pdf. Such data can be for instance from genetic origin (such as mutations) or from other variables with slow characteristic evolution time. We stress that it is the probability distribution of the unmodelables that has to be close to equilibrium: note that this does not require the value of the unmodelable variables to reach a constant value but the values should be drawn from a steady state distribution.

We see room for improvement also concerning the selection of the density estimation method. We adopted a well known form of nonparametric estimation through kernel density estimation, but other approaches could be tailored to a specific problem - especially when high-dimensional datasets come into play [48]. Moreover, introducing density estimation methods able to integrate categorical variables would greatly benefit the technique, especially in biomedical problems (e.g. it would be extremely beneficial to include the grade of a tumor, or the particular sequence of therapies that a patient has undergone). The modularity of the BaM^3^ method makes it extremely versatile, allowing one to change the density estimation step, the modeling part or both of them at the same time to improve the final prediction scores.

Care should also be taken to generate pdfs that are able to cope with outliers. In our proof of concept we generated the probability distributions considering the same weight for every patient, irrespective of his position in the parameter space. Techniques able to identify these extreme cases and to improve their contribution to the final pdfs should be implemented for a better method performance [49].

In summary, we can identify three main actions that could be undertaken when these limitations hamper the predictive capabilities of BaM^3^: (i) one should look for ways to improve the mathematical model, designing it to be as informative as possible; (ii) then, an effort should be put to constrain the model by a robust choice of parameters; finally, (iii) an extreme care should be devoted to the selection of the most informative unmodelable variables.

We conclude by stating that the proposed method is not restricted to oncology. The core problem concerning clinical predictions is that data are heterogeneous and sparse in time along with lack of full mechanistic knowledge. Therefore, a vast variety of medical problems could be addressed by using the BaM^3^ approach. For instance, predicting the fate of renal grafts by using pre- and post transplantation data is a prime application of our proposed methodology.

## Methods

### Formal definition of BaM^3^

We start by assuming a random variable (r.v.) triplet (**Y**, **X**_*m*_, **X**_*u*_) that denotes the system’s modelable **X**_*m*_, unmodelable **X**_*u*_ variables/data (e.g. patient’s age or sex, results of different ‘-omics’ techniques, etc.) and the associated observed clinical outputs **Y**. We then introduce *t*_0_ as the clinical presentation time of a patient at which the patient specific r.v. realizations 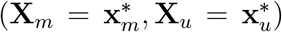 are obtained. The overall goal of the method is to predict the patient’s clinical outputs by an estimate 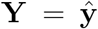 at a certain prediction time *t_p_*. The true clinical outputs of the patient will be denoted as **y**. Moreover, we consider the existence of a *N –*patient ensemble dataset (**y**, **x**_*m*_, **x**_*u*_). In this dataset all the variables (i.e. modelables, unmodelables and clinical outputs) are recorded at the time of diagnosis *t_d_*, which might differ from one patient to another. Both *t*_0_ and *t_d_* are calculated from the onset of the disease. We introduce two distinct times to account for the variability of the disease stage among different patients (*t_d_*) and the time at which a specific patient is presented to the clinic (*t*_0_) (see the corresponding SI Fig. S15).

The core idea of the method is to consider the predictions of the mathematical model 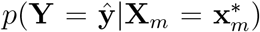 as an informative Bayesian prior of the posterior distribution 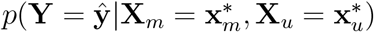. We can prove that:

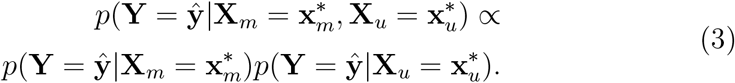

The implementation of the BaM^3^ method therefore reduces to the calculation of the aforementioned probability distributions. Although the prediction of the probability distribution function (pdf) of the clinical outputs is rather straightforward for the mathematical model (see the following Methods sections), obtaining the pdf of the patient’s unmodelable data is not trivial. To retrieve the latter, we use a density estimator method upon the patient ensemble dataset to derive *p*(**Y**, **X**_*u*_), and then consider the patient specific realization 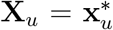. For further details about method derivation and estimators of performance see the corresponding SI sections.

### Testing the method on synthetic glioma growth

The equations of the selected mathematical model [17, 18] (‘full model’) describe the spatio-temporal dynamics of tumor cell density (*c*), oxygen concentration (*n*) and vascular density (*v*) in the context of glioma tumor growth. The full model includes the variation of cell motility and proliferation due to phenotypic plasticity of tumor cells induced by microenvironmental hypoxia [15, 50, 51, 52, 53]. It also accounts for oxygen consumption by tumor cells, formation of new vessels due to tumor angiogenesis and vaso-occlusion by compression from tumor cells [54, 55, 18]. We generate *N* = 500 virtual patients by sampling the parameters of the full model from a uniform distribution over the available experimental range. We consider the tumor cell spatial density *c* to be the modelable variable. Moreover, we treat the integral over the tissue of oxygen concentration and vascular density, denoted as 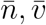 respectively, as the unmodelable quantities. Starting from the same initial conditions, we simulate the behavior of each virtual patient for three years, storing the values of all variables at each month.

As sketched in Fig. 8, we use the modelable variable to setup a mathematical model. In particular, we take *c*(*x, t*_0_) at a specific time point, the clinical presentation time *t*_0_, and use it as the initial condition for a Fisher–Kolmogorov equation [22, 56, 57, 58, 59] (‘FK model’). We use this model to predict tumor behavior at a specific time in the future, the prediction time *t_p_*. For each simulated patient we calculate the Tumor Size (TS) and Infiltration Width (IW). In parallel, for each patient we evaluate the diagnosis time *t_d_* as a random number in the interval [*t*_0_ − 6, *t*_0_ + 6] (in the unit of months), and collect the values of modelables, unmodelables and clinical outputs at this time to build the patient ensemble. Given the patient-specific modelable and unmodelable variables (*c*(*x, t*_0_) and 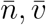, respectively) at the clinical presentation time *t*_0_, the BaM^3^ method therefore produces the probability of observing the TS and IW at a specific prediction time *t_p_*.

**Figure 8:**
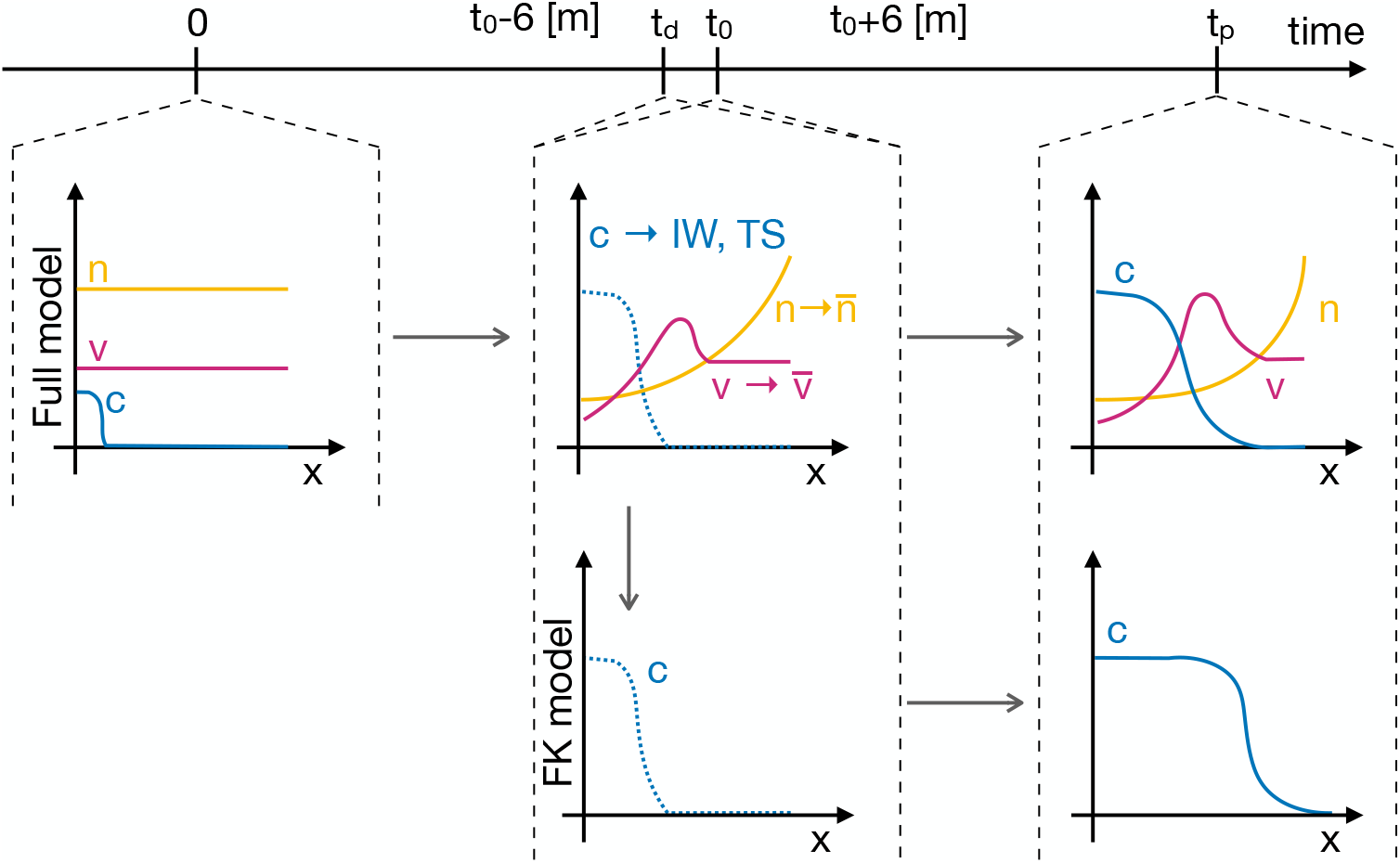
Workflow for generating the synthetic data. The full model is initialized at *t* = 0 and used to simulate the spatio-temporal variation of tumor density *c*, oxygen concentration *n* and functional tumor-associated vasculature *v*. For each virtual patient, two clinical outputs are tracked, i.e. the Infiltration Width (IW) and Tumor Size (TS), and two unmodelables recorded, i.e. the oxygen and vasculature integral over the tissue 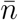 and 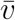, respectively. These quantities are generated at the time of clinical presentation *t*_0_ every time the method is applied. To generate the patient ensemble, they are also generated at the diagnosis time *t_d_*. The latter is assumed to be a random time between *t*_0_ ± 6 months. Then, the spatial profile of tumor concentration at time *t*_0_ is used as the initial condition for the Fisher–Kolmogorov (FK) model, which is in turn used to simulate tumor growth until the prediction time *t_p_*.

### Mathematical models for glioma growth

The system variables are the density of glioma cells *c*(*x, t*), the concentration of oxygen *n*(*x, t*) and the density of functional vasculature *v*(*x, t*) [17, 18]. For simplicity we consider a one-dimensional computational domain. We normalize the system variables to their carrying capacity and write the system as

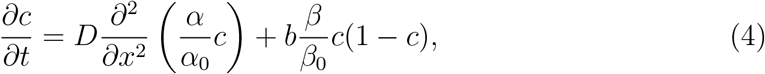

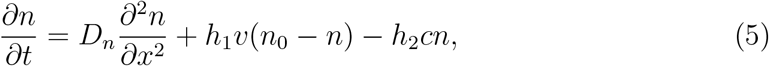

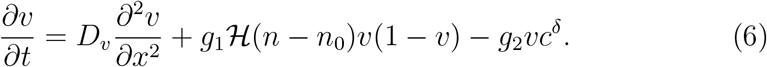

Here 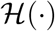 is a sigmoidal function 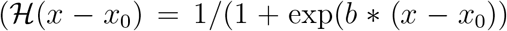, with *b* > 0 being a constant) allowing for tumor angiogenesis in hypoxic conditions, i.e. for *n < n*_0_ where *n*_0_ is the hypoxic oxygen threshold. Then, the functions *α* = *α*(*n*) and *β* = *β*(*n*) account for the dependence of cellular motility and proliferation on the oxygen level, respectively [15, 50, 52]. They are defined as:

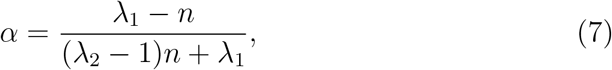

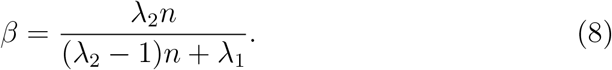

When the oxygen level is fixed to the maximum level *n* = 1 in the tissue *α* = *α*_0_ and *β* = *β*_0_, so that the equation for *c* reduces to

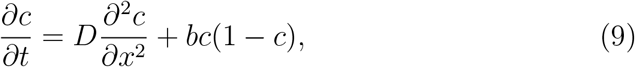

which we denote in the rest of the manuscript as the Fisher-Kolmogorov (FK) model for tumor cell density. We remark that equation (9) has been extensively used to predict untreated glioma kinetics based on patient-specific parameters from standard medical imaging procedures [22, 56, 57, 58, 59].

Eq. (4)-(6) define an extended version of the FK equation, enriched with nonlinear glioma cell diffusion and proliferation terms. The latter terms depend on the oxygen concentration in the tumour microenvironment, which is in turn coupled to cell density through the oxygen consumption term. The functional vascular density controls the supply of oxygen to the tissue. Blood vessel density increases due to tumor angiogenesis and decreases because of vaso-occlusion by high tumor cell density. The values of the parameters used in the simulations and their descriptions are given in Tab. S1. In addition, a typical full model simulation is shown in Fig. S2 for a representative patient.

We solve the system in (4)-(6) by imposing the initial conditions:

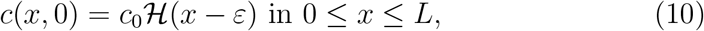

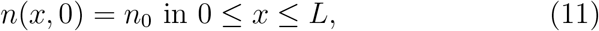

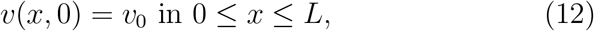

where the positive parameters *c*_0_, *n*_0_ and *v*_0_ are the initial density of glioma cells spatially distributed in a segment of length *ε*, the density of functional tumour vasculature and the oxygen concentration, respectively. Then, *L* > 0 is the length of the one-dimensional computational domain. In addition, we consider an isolated host tissue in which all system behaviours arise solely due to the interaction terms in Eq. (4)-(6). This assumption results in no-flux boundary conditions of the form:

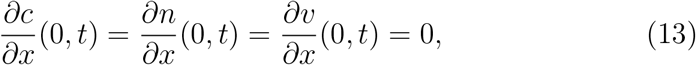

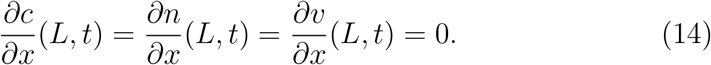

Both the full and FK models are used to calculate two clinical outputs, namely the tumor Infiltration Width (IW) and Tumor Size (TS). The IW at a specific time is defined by the difference between the points where glioma cell density is 80% and 2% of the maximum cellular density. In turn, the TS is obtained by integrating the spatial profile of tumor density and dividing it for the maximum value of the latter.

We run the full model and simulate the growth of the tumor for *N* patients, each one from a parameter set taken randomly from a uniform distribution over the parameter range. We run simulations for *N* =50,100,250 and 500 with 10 repetitions within each N-case. To generate the patients, we vary five parameters in the list in Tab. S1, namely the tumor motility *D*, proliferation rate *b*, oxygen consumption *h*_1_, vascular formation and occlusion rates *g*_1_ and *g*_2_, respectively. Then, we use the tumor density at the time of clinical presentation, *t*_0_, as the initial condition for the FK model. The latter model is employed to generate predictions at the prediction time *t_p_*. We also consider the unmodelable variables and clinical outputs at the diagnosis time *t_d_*, taken randomly between *t*_0_ ±6 months, to build the patient ensemble. Finally, we use the results of the full model in terms of clinical outputs as the ground truth to be compared with the predictions of the FK model alone and with the ones obtained by the BaM^3^ method.

### Probability distribution from the FK model

As described in the previous sections, we take the spatial profile of tumor density at the clinical presentation time *t*_0_ as the initial condition of the FK model. We use the latter mathematical model to run simulations over the whole parameter set for cell motility *D* and proliferation rate *b*. Then, we define the model-derived pdf as in the following. For each couple of clinical outputs IW^∗^ and TS^∗^ we calculate the area *A_α_*(IW^∗^,TS^∗^) over the (IW,TS) plane as *A_α_* = [(1 − *α*)IW^∗^ < IW < (1 + *α*)IW^∗^), (1 − *α*)TS^∗^ < TS < (1 + *α*)TS^∗^)], where *α* is a given tolerance (here set equal to *α* = 0.05). Then, we calculate the pdf by normalizing *A_α_* by the total area of predicted IW and TS values. We store the value of the probability for each patient at the different prediction times and use it to compute the expected value of the model pdf.

### Probability distribution of the unmodelables from the full model

To retrieve the data-derived pdf we use a normal kernel density estimator (KDE) [19, 20] which depends upon all the data points in the patient ensemble. Briefly, the method estimates the joint probability 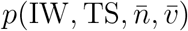 from which the ensemble entries are drawn through the sum of a kernel function over all the occurrences of the dataset. The kernel function is characterized by a hyper-parameter, the bandwidth 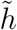, which we assume according to Silverman’s rule of thumb

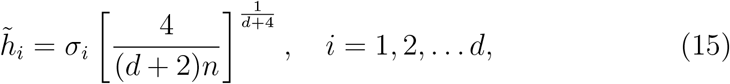

where *d* is the number of dimensions, *n* is the number of observations, and *σ_i_* is the standard deviation of the i-th variate [60]. After calculating 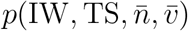, we specify the realization of a specific patient and calculate the value of 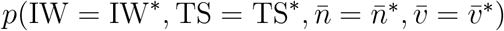 over the (IW,TS) space of the estimated clinical outputs.

### Scoring glioma growth predictions

We calculate for each patient the relative errors *d_m_* and *d_b_* as described the main text. To assess how the BaM^3^ method has changed the prediction of the mathematical model we compare the latter quantities: if |*d_b_* − *d_m_*| ≤ *εd_m_*, then there was no change; if *d_b_* > (1 + *ε*)*d_m_*, then the method deteriorated the prediction of the model; if *d_b_* < (1 − *ε*)*d_m_*, then the method improved the prediction of the model. Here, *ε* is a tolerance used for the comparison, taken to be equal to *ε* = 0.05.

### Calculation of the effective variance

To calculate the effective variance *s* shown in Fig. 3 we first calculate the mixed central moments Σ_*ij*_ of the pdf of interest according to the formula

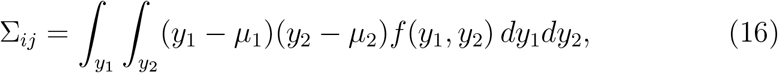

where *y*_1_ and *y*_2_ are the clinical outputs (IW and TS, respectively) and *μ*_1_, *μ*_2_ the expected values of the corresponding variables. The elements of Σ form a symmetric two-dimensional matrix, for which we calculate the determinant. We define the effective variance *s* as the natural logarithm of the latter determinant. In Eq. (16) we consider *f*(*y*_1_*, y*_2_) to be the pdf from the mathematical model or from the BaM^3^ method depending on whether we are interested in the effective variance *s_m_* or *s_b_*, respectively.

### Pdf from the two-compartment model in CLL

Messmer and colleagues [28] measured the fraction of labelled B-CLL cells in a court of 17 CLL patients that were administered deuterated water. They calibrated a two-compartment model on each patient and were able to reproduce the kinetics of labeled cells over a long time. We adopt their model and use it to generate the pdf for the CLL example. The fraction of labeled cells over time is calculated through the expression

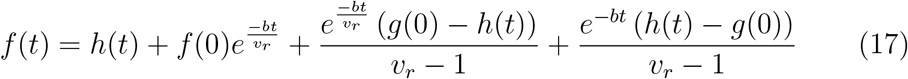

where *g*(0) is the initial fraction of cells in the first compartment, *b* the fractional cell birth, *v_r_* the relative size of the compartments and *h*(*t*) is the deuterated water concentration of the body over time. The latter is a function of the fractional daily water exchange *f_w_*. We refer the interested reader to the supplementary information of Messmer et al. [28] for a more detailed description of the model and a full account of the model parameters. In this work we focus on three quantities, namely *b*, *v_r_* and *f_w_*, and run the model in Eq. (17) over the experimental range. This range was obtained by considering the patient-specific fitting performed by Messmer and colleagues and selecting the minum and maximum values. We evaluate the fraction of labeled cells at day 50, *f*_50_, and build the probability distribution from its histogram, by counting the number of occurrences of a given 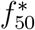 for 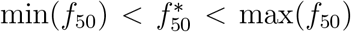 and then normalizing the result. For the CLL example all the patients start with the same initial fraction of labeled cells, set to zero.

### Pdf from the patients’ unmodelables in CLL

The data-derived pdf in the CLL example is obtained from four unmodelable quantities that are measured for each patient during the study. We consider all the possible combinations of unmodelables and calculate the MSE for each case, as shown in the inset of Fig. 6. The scatter plot in the same picture refers to the case in which the CD38 expression (*x*_*u*,1_), age (*x*_*u*,2_), growth rate of white blood cells (*x*_*u*,3_) and V_H_ mutation status (*x*_*u*,4_) are added consecutively with the specified order. As in the glioma example we build the sub-dataset (*y*, **x**_*u*_), where *y* and **x**_*u*_ = (*x_u,i_*) are the *f*_50_ and unmodelable variables of each patient respectively, and apply the KDE using Silverman’s rule for the hyperparameters. The requested pdf, i.e. 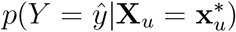 is obtained by conditioning the probability from the KDE with the realizations of the unmodelables of the specific patient and calculating the result over the range of the estimated clinical output 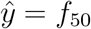.

### Mathematical model for ovarian cancer

We assume the total number of tumor cells *T* to be composed of the sensitive *S* and resistant *R* subpopulations. The latter are described by the following system of ODEs:

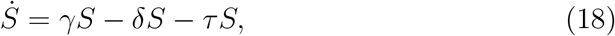

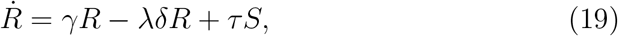

where *γ* is the tumor net growth rate, *δ* = *δ*(*t*) is the death rate induced by chemotherapy, *τ* is the mutation rate from sensitive to resistant cells, and *λ* is a factor that accounts for reduced death by therapy in resistant cells. As detailed in Fig. S11, the treatment is composed of three phases: first, the patients undergo different cycles of NACT; then, surgery is performed. The latter reduces the total tumor volume, irrespective of cells being sensitive or resistant, of a factor *β*. Finally, another series of chemotherapy cycles are performed. During chemotherapy, *δ* = *δ*_0_, whereas we set this parameter to zero after chemotherapy and until tumor relapse. The latter condition occurs when *T* reaches the value *T_R_*.

Equations (18)-(19) can be analytically integrated, and their results used to build the probability distribution of the clinical output - TtR, in this case. To obtain the pdf from the model we calculate the time the tumor takes to reach the cell number at relapse *T_R_* starting from the cell number after therapy. We perform this calculation using the initial tumor cell number of each patient, and by varying both the initial fraction of *S* cells, *x*_0_, and the chemotherapy-induced death rate, *δ*_0_. We then obtain the patient-specific probability distribution from the histogram of TtR, similarly to what done in the CLL section. For *x*_0_ we select a range between 0.4 and 0.9, accounting for tumors with different initial degrees of intrinsic resistance [36]. For *δ*_0_, we first use a uniform distribution between 0.1 and 10 days^−1^, accounting for a wide variation in death rates. The latter choice produces an almost flat distribution for the clinical output (see Fig. S12). To improve the mathematical model parametrization, we use the information about the tumor volume change after the first cycle of chemotherapy, which is included in the dataset. By fitting *T* obtained from (18)-(19) to the observed volume change, we find a value of *δ*_0_ for each patient in the dataset [36]. We take the mean value of these rates and use it to update the model pdf (see Fig. S13). We consider a range for *δ*_0_ which is centered around its mean value across the patients, within an interval of ± 40%. Selecting other ranges provides similar results, however a variation of 40% returns the lowest MSE. Analytical integration of (18)-(19), as well as additional details about model parametrization are available in the Supplementary Material.

### Unmodelable variable for the ovarian cancer study

We build the data-derived pdf for the ovarian cancer example by exploiting the information about the age of the patients at diagnosis. Similarly to what done in the previous test cases, we first build the sub-dataset (TtR, *A*) by entering the information of each patient (here, *A* is the patient age). Then, we apply the KDE using Silverman’s rule to estimate the bandwidth and calculate the joint probability *p*(TtR*, A*). The data-derived pdf for each patient *p*(TtR|*A*^∗^) is finally obtained over the domain of the clinical output TtR by considering the patient-specific age *A* = *A*^∗^.

## Data availability

The data and code used to generate the simulations and the plots is available at https://github.com/PiMasGit/BaM3-method

## Author contributions

Conceptualization, H.H.; study design, P.M., S.S., J.C.L.A., H.H.; software, P.M., S.S., J.C.L.A.; formal analysis, P.M., S.S, J.C.L.A., M.M.-H., H.H.; writing—original draft preparation, P.M.; writing—review and editing, P.M., S.S., J.C.L.A., M.M.-H., H.H.; supervision, M.M.-H., H.H.; funding acquisition, M.M.-H., H.H.

## Acknowledgments

The authors gratefully acknowledge the funding support of the Helmholtz Association of German Research Centers—Initiative and Networking Fund for the project on Reduced Complexity Models (ZT-I-0010). HH and PM acknowledge the funding support of MicMode-I2T (01ZX1710B) by the Federal Ministry of Education and Research (BMBF). H.H. is supported by SYSIMIT (01ZX1308D) and MulticellML (01ZX1707C) by the Federal Ministry of Education and Research (BMBF). Finally, HH would like to thank the Volkswagenstiftung for the its support within the “Life?” programm (96732).

