## Supplementary Information for "Bayesian combination of mechanistic modeling and machine learning (BaM^3^): improving personalized tumor growth predictions"

April 21, 2021

### Derivation of the method

To derive equation Eq. (3) in the main text, we consider the joint probability $p(\mathbf{Y}, \mathbf{X}_m, \mathbf{X}_u)$. This can be written in two different ways:

$$p(\mathbf{Y}, \mathbf{X}_m, \mathbf{X}_u) = p(\mathbf{Y}|\mathbf{X}_m, \mathbf{X}_u)p(\mathbf{X}_m, \mathbf{X}_u), \quad (1)$$
$$p(\mathbf{Y}, \mathbf{X}_m, \mathbf{X}_u) = p(\mathbf{X}_m, \mathbf{X}_u|\mathbf{Y})p(\mathbf{Y}). \quad (2)$$

If we assume statistical independence between the conditioned modelable and unmodelable variables the latter equation becomes

$$p(\mathbf{Y}, \mathbf{X}_m, \mathbf{X}_u) = p(\mathbf{X}_m|\mathbf{Y})p(\mathbf{X}_u|\mathbf{Y})p(\mathbf{Y}), \quad (3)$$

*To whom correspondence should be addressed.


and, after applying Bayes theorem we obtain

$$p(\mathbf{X}_m|\mathbf{Y})p(\mathbf{X}_u|\mathbf{Y})p(\mathbf{Y}) = p(\mathbf{X}_u|\mathbf{Y})p(\mathbf{Y}|\mathbf{X}_m)p(\mathbf{X}_m) = \quad (4)$$

$$= \frac{p(\mathbf{Y}|\mathbf{X}_u)p(\mathbf{X}_u)}{p(\mathbf{Y})}p(\mathbf{Y}|\mathbf{X}_m)p(\mathbf{X}_m) = \quad (5)$$

$$= p(\mathbf{Y}|\mathbf{X}_u)p(\mathbf{Y}|\mathbf{X}_m)\frac{p(\mathbf{X}_u)p(\mathbf{X}_m)}{p(\mathbf{Y})}. \quad (6)$$

Then, we equate the rhs of equations (1) and (6) and obtain the relationship:

$$p(\mathbf{Y}|\mathbf{X}_m, \mathbf{X}_u) = \frac{p(\mathbf{Y}|\mathbf{X}_u)p(\mathbf{Y}|\mathbf{X}_m)}{p(\mathbf{Y})}\frac{p(\mathbf{X}_m)p(\mathbf{X}_u)}{p(\mathbf{X}_m, \mathbf{X}_u)}. \quad (7)$$

Considering the realization of a specific patient $(\mathbf{x}_m^*, \mathbf{x}_u^*)$, we recover Eq. (3) of the main text:

$$p(\mathbf{Y} = \hat{\mathbf{y}}|\mathbf{X}_m = \mathbf{x}_m^*, \mathbf{X}_u = \mathbf{x}_u^*) \propto p(\mathbf{Y} = \hat{\mathbf{y}}|\mathbf{X}_m = \mathbf{x}_m^*)p(\mathbf{Y} = \hat{\mathbf{y}}|\mathbf{X}_u = \mathbf{x}_u^*). \quad (8)$$

We remark that the first term on the rhs of the latter equation is the model-derived pdf, whereas the second term is the data-driven distribution. Both of them contribute to the final prediction, in a multiplicative way under our assumptions.

Finally, we state the previously derived findings in terms of entropies. This allows for a graphical representation of the method structure in terms of a Venn diagram. For this, we apply the logarithm to both sides of Eq. (7) and obtain:

$$\log[p(\mathbf{Y}|\mathbf{X}_m, \mathbf{X}_u)] = \log[p(\mathbf{Y}|\mathbf{X}_m)] + \log[p(\mathbf{Y}|\mathbf{X}_u)] - \log[p(\mathbf{Y})] - \log[\frac{p(\mathbf{X}_m, \mathbf{X}_u)}{p(\mathbf{X}_m)p(\mathbf{X}_u)}], \quad (9)$$

which, after multiplication for the joint probability $p(\mathbf{X}_m, \mathbf{X}_u, \mathbf{Y})$ and integration over the random variable space gives

$$S(\mathbf{Y}|\mathbf{X}_m, \mathbf{X}_u) = S(\mathbf{Y}|\mathbf{X}_m) + S(\mathbf{Y}|\mathbf{X}_u) - S(\mathbf{Y}) + I(\mathbf{X}_m, \mathbf{X}_u). \quad (10)$$

Eq. (10) can be translated to a Venn diagram for graphical representation. Fig. S1**A** shows the diagram when the probabilities $p(\mathbf{X}_m|\mathbf{Y})$ and $p(\mathbf{X}_u|\mathbf{Y})$ are assumed to be independent, as in our proof. When this assumption does not hold, the mutual information $I(\mathbf{X}_m, \mathbf{X}_u)$ does not coincide with $I(\mathbf{X}_m, \mathbf{X}_u, \mathbf{Y})$, but an additional term, i.e. $I(\mathbf{Y}|\mathbf{X}_m, \mathbf{X}_u)$, should be subtracted from the latter (Fig. S1**B**).

#### Performance estimators of the method: a Cramer-Rao bound

Here we would like to point out that the only assumption made to derive expression (8) was the statistical independence of the modelable and unmodelable variables conditioned by the clinical outputs, i.e. :

$$p(\mathbf{X}_m, \mathbf{X}_u|\mathbf{Y}) = p(\mathbf{X}_m|\mathbf{Y})p(\mathbf{X}_u|\mathbf{Y}). \quad (11)$$

The impact of this assumption can be estimated by reformulating (8) into the corresponding information theoretical quantities. Then, it can be proven that

$$S_{method}(\mathbf{Y}|\mathbf{X}_m, \mathbf{X}_u) \geq S_{true}(\mathbf{Y}|\mathbf{X}_m, \mathbf{X}_u). \tag{12}$$

This highlights that the entropy $S_{method}$ produced by the proposed BaM$^3$ method is greater than the entropy $S_{true}$ of the corresponding 'true' predictions without using the assumption (11). In particular, we can use the inequality that connects the mean square estimator (MSE) between reality and BaM$^3$ prediction and the conditional entropy that reads [1]:

$$\langle(\hat{\mathbf{y}}(\mathbf{x_m}, \mathbf{x_u}) - \mathbf{y})^2\rangle \geq \frac{e^{S_{method}(\mathbf{Y}|\mathbf{X}_m,\mathbf{X}_u)}}{2\pi e} \geq \frac{e^{S_{true}(\mathbf{Y}|\mathbf{X}_m,\mathbf{X}_u)}}{2\pi e}. \tag{13}$$

The above inequality implies that the assumption of statistical independence of the two sets of variables forces the lower bound of the MSE higher in comparison to a method that would not assume this. The above inequality is a generalization of the Cramer-Rao bound, which can be recovered by assuming a Gaussian distribution for the r.v. $(\mathbf{Y}|\mathbf{X}_m, \mathbf{X}_u)$.

#### Rationale

We would like to provide some arguments on our rational, i.e. why BaM$^3$ improves prediction performance. As a cost function that quantifies prediction performance, we consider the mean square error (MSE) given by $(\hat{\mathbf{y}} - \mathbf{y})^2$. As stated above, the main idea is to use the mechanistic model predictions as an informative prior. In turn, the unmodelable data should contribute to the corresponding likelihood term. The corresponding Bayesian formula, after the assumption (3), leads to Eq. (8) which can be alternatively written as:

$$p(\mathbf{Y}|\mathbf{X}_m, \mathbf{X}_u) \propto p_{unmodel} \times p_{model}, \tag{14}$$

where the prior is $p_{model} = p(\mathbf{Y}|\mathbf{X}_m)$, and the corresponding unmodelable-based likelihood is $p_{unmodel} = p(\mathbf{Y}|\mathbf{X}_u)$.

To illustrate the potential of the proposed BaM$^3$ method, let's assume an estimator of clinical outputs based on the full input dataset $(\mathbf{x}_m^*, \mathbf{x}_u^*)$, e.g. using a certain machine learning method. Any estimator is dictated by the Cramer-Rao inequality [2] which provides a lower bound for the MSE:

$$\langle(\hat{\mathbf{y}} - \mathbf{y})^2\rangle \geq \frac{1}{\mathrm{I}_{full}}, \tag{15}$$

where $\mathrm{I}_{full} = \mathrm{I}(\hat{\mathbf{y}})$ is the corresponding Fisher Information (FI) of the full dataset estimator. Thus, even for unbiased estimators this lower bound cannot be less than inverse of FI which is always a finite number.

In the BaM$^3$ method, we can improve the above lower bound by adding an informative prior based on mathematical modeling. In particular, by using the van Trees inequality [3], known as Bayesian Cramer-Rao bound, and assuming an increasing model-based FI denoted as $\mathrm{I}_{model}$, we can deduce that the lower bound vanishes:

$$\langle (\hat{\mathbf{y}} - \mathbf{y})^2 \rangle \geq \lim_{\mathrm{I}_{model} \to \infty} \frac{1}{\mathrm{I}_{umodel} + \mathrm{I}_{model}} = 0. \tag{16}$$

The expression $\mathrm{I}_{umodel} \leq \mathrm{I}_{full}$ denotes the FI that unmodelable data achieve. The last formula shows that the proposed BaM$^3$ method has the potential to minimize the MSE lower bound and outscore any data-driven method by harnessing the benefits of a "good" mathematical model.

### Mathematical models for ovarian cancer

#### Model with two cell subpopulations

We model the tumor cell number $T$ as the sum of sensitive $S$ and resistant $R$ cells. These two subpopulations are described by the system of ODE:

$$\dot{S} = \gamma S - \delta S - \tau S, \tag{17}$$

$$\dot{R} = \gamma R - \lambda \delta R + \tau S, \tag{18}$$

where $\gamma$ is the tumor net growth rate, $\delta = \delta(t)$ is the death rate induced by chemotherapy, $\tau$ is the mutation rate from sensitive to resistant cells, and $\lambda$ is a factor that accounts for reduced death by therapy in resistant cells. During chemotherapy we assume $\delta = \delta_0$, otherwise this parameter is set to zero (see Figure S11). Since equation (17) does not depend on $R$, we first analytically integrate it and then substitute the value of $S(t)$ in (18). This allows to also solve the equation for $R$. Thus, we can write the solution for $S$ and $R$ as

$$S(t) = S_0 e^{(\gamma - \delta_0 - \tau)t}, \tag{19}$$

$$R(t) = \left[ R_0 + \frac{\tau S_0}{\delta_0(1-\lambda) + \tau} \right] e^{(\gamma - \lambda\delta_0)t} - \frac{\tau S_0}{\delta_0(1-\lambda) + \tau} e^{(\gamma - \delta_0 - \tau)t}, \tag{20}$$

for $0 < t < t_1$, where $t_1$ is the end of the first treatment stage with chemotherapy. The initial number of sensitive and resistant cells are given by $S_0 = x_0 T_0$ and $R_0 = (1 - x_0) T_0$, respectively, where $x_0$ and $T_0$ are the initial fraction of sensitive cells in the total population and the initial number of tumor cells.

After the first round of chemotherapy, a surgery is performed to remove a constant fraction $\beta$ of the tumor. We write the number of cells after surgery as

$$S_2 = \beta S_1, \tag{21}$$

$$R_2 = \beta R_1, \tag{22}$$

where $S_1$ and $R_1$ are the values of $S$ and $R$ at $t = t_1$.

Finally, we use $S_2$ and $R_2$ after surgery to build the analytical solution of the system for $t_1 < t < t_d$, over the duration of the second stage of chemotherapy (see the scheme in Figure SX). For this time frame, $S$ and $R$ are given by

$$S(t) = S_2 e^{(\gamma-\delta_0-\tau)(t-t_1)}, \tag{23}$$

$$R(t) = \left[R_2 + \frac{\tau S_2}{\delta_0(1-\lambda)+\tau}\right] e^{(\gamma-\lambda\delta_0)(t-t_1)} - \frac{\tau S_2}{\delta_0(1-\lambda)+\tau} e^{(\gamma-\delta_0-\tau)(t-t_1)}. \tag{24}$$

After the last cycle of chemotherapy the tumor resumes exponential growth, at the net growth rate $\gamma$. The time-to-relapse (TtR) is calculated as the time the tumor takes to reach the relapse cell number $T_R$ starting from the cell number after therapy, $T_d = S_d + R_d$, where $S_d$ and $R_d$ are obtained from (23)-(24) at $t = t_d$. Therefore, the following relation holds for the clinical output TtR:

$$\text{TtR} = \frac{1}{\gamma} \ln\left(\frac{T_R}{T_d}\right). \tag{25}$$

The parameters that enter the model are listed in Table S2. Most of them are taken from the publication from which we obtained the patient clinical data [4]. For each patient, the dataset reports the initial tumor cell number, the cell number after the first stage of chemotherapy, and the age. As detailed in the Methods section in the main text, the age variable is used to generate the data-driven pdf in the context of the BaM$^3$ methodology. We use the change in tumor size after the first round of chemotherapy to calculate $\delta_0$ for each patient, by means of equations (19)-(20) calculated at $t = t_1$. Then we calculate the mean of all these values and use it to parametrize the mathematical model and, together with varying the initial fraction of sensitive cells $x_0$, generate the corresponding probability distribution. This version of the model, denoted as 'fitted', is compared to its 'uninformative' counterpart, in which we assume an arbitrary range for the death rate $\delta_0$. The pdf obtained in the two cases are displayed in Figures S12 and S13.

#### Simplified mathematical model

In addition to the two population framework of the previous section, we implemented a simplified version of the mathematical model. We consider a single cell population and an effective death rate $\delta_e$ that is enforced over all the duration of the therapy, taking into account both rounds of chemotherapies and surgery. The tumor cell number varies in time according to

$$T = \begin{cases} T_0 e^{(\gamma-\delta_e)t}, & \text{for } 0 < t < t_d, \\ T_d e^{\gamma(t-t_d)}, & \text{for } t_d < t < t_R, \end{cases} \tag{26}$$

where $T_0$, $T_d$, $\gamma$, $t_d$ and $t_R$ are the initial and after therapy tumor cell number, net growth rate, duration of therapy and time of tumor relapse, respectively. In this setting, the time-to-relapse TtR is obtained via equation

$$\text{TtR} = \frac{1}{\gamma} \ln\left(\frac{T_R}{T_0}\right) - \frac{\gamma-\delta_e}{\gamma} t_d, \tag{27}$$

where $T_R$ is the tumor cell number at relapse. We apply the BaM$^3$ framework as done in the previous section using this simplified model instead of the two population version. The initial tumor cell number is available for each patient, whereas we vary only the effective death rate $\delta_e$ to generate the model pdf. To define a parameter range for $\delta_e$, we first fit this quantity for each patient by making use of Equation (26) at $t = t_1$ and the available patient-specific data about tumor cell number at this time (similarly to what done for the 'fitted' case of the two populations model in the previous section). Then, we take the mean of $\delta_e$ over all patients and define the interval of this parameter to enclose values below and above 50% of the mean value. Results for this modeling choice are available in Figure S14, which shows the pdf obtained from the simplified mathematical model, density estimation and BaM$^3$ approach. Note that for the density estimation we adopted the same unmodelable as in the previous section for comparison purposes. The Figure shows the poor performance of the BaM$^3$ method when a poor modeling strategy is enforced. In this case, the model is not able to provide a suitable prior to the data-driven pdf, which fails to be improved in many occurrences. Indeed, the MSE after applying BaM$^3$ using the simplified model amounts to MSE = 50.513 months$^2$, which significantly higher compared to the MSE from the 'fitted' two populations case (MSE = 30.895 months$^2$).

Table S1: Values and description of the parameters for the full model. The reference sources for the parameters are available in [5].

| Parameter | Description | Value |
|---|---|---|
| $D$ | Intrinsic diffusion rate of glioma cells | $[2.73 \times 10^{-3}, 2.73 \times 10^{-1}]\mathrm{mm}^2\mathrm{d}^{-1}$ |
| $b$ | Intrinsic proliferation rate of glioma cells | $[2.73 \times 10^{-4}, 2.73 \times 10^{-2}]\mathrm{d}^{-1}$ |
| $\lambda_1$ | Phenotypic switching parameter | 2 |
| $\lambda_2$ | Phenotypic switching parameter | 1 |
| $D_n$ | Diffusion rate of oxygen | $1.51 \times 10^{2}\mathrm{mm}^2\mathrm{d}^{-1}$ |
| $h_1$ | Oxygen supply rate | $3.37 \times 10^{-1}\mathrm{d}^{-1}$ |
| $h_2$ | Oxygen consumption rate | $[5.73 \times 10^{-1}, 1.14 \times 10^{1}]\mathrm{d}^{-1}$ |
| $D_v$ | Vasculature dispersal rate | $5 \times 10^{-4}\mathrm{mm}^2\mathrm{d}^{-1}$ |
| $g_1$ | Vasculature formation rate | $[10^{-2}, 2.5 \times 10^{-1}]\mathrm{d}^{-1}$ |
| $g_2$ | Vasculature occlusion rate | $[5.0 \times 10^{-1}, 1.5 \times 10^{1}]\mathrm{d}^{-1}$ |

Table S2: Values and description of the parameters for the ovarian cancer study. The reference sources for the parameters are available in [4].

| Parameter | Description | Value |
|---|---|---|
| $\gamma$ | Tumor net growth rate | $5.8 \times 10^{-3}\mathrm{d}^{-1}$ |
| $\beta$ | Tumor reduction by surgery | 0.01 |
| $\lambda$ | Therapy reduction factor for resistance cells | 0.01 |
| $\tau$ | Mutation rate from sensitive to resistance cells | $1.6 \times 10^{-5}\mathrm{d}^{-1}$ |
| $T_R$ | Tumor cell number at relapse | $10^9$ |
| $t_1$ | Duration of first round of chemotherapy | 63d |
| $t_d$ | Total duration of therapy | 126d |

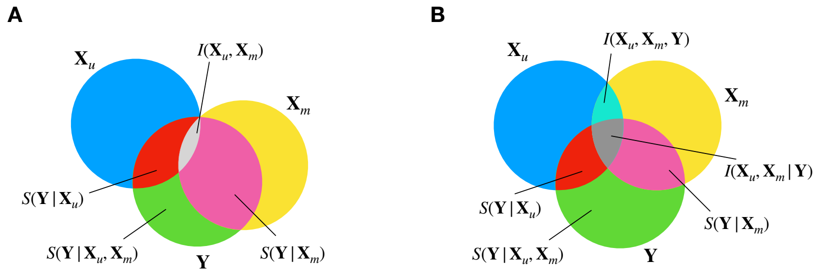


Figure S1: Venn diagrams representing the relationships between the quantities that appear in the method. In particular, the cases for which the probabilities $p(\mathbf{X}_m|\mathbf{Y})$ and $p(\mathbf{X}_u|\mathbf{Y})$ are independent (**A**) or not (**B**) are shown.

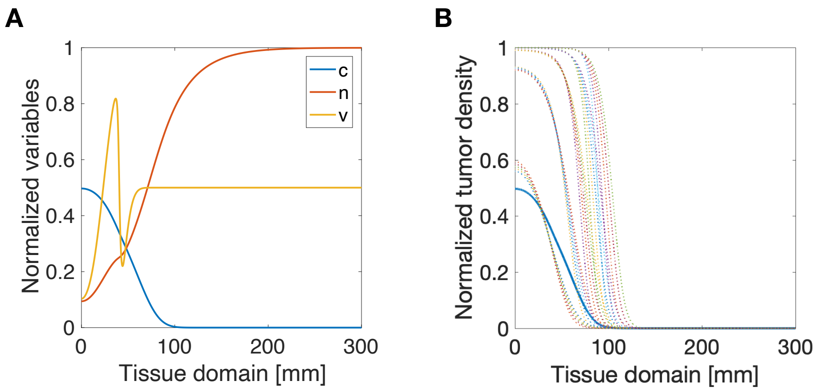


Figure S2: **A** Spatial profile of the normalized variables simulated by the full model. The plot is for a representative patient after 3 years from the beginning of the simulation. Here, $c$, $n$ and $v$ are the normalized tumor cell density, oxygen and vascular density, respectively. **B** Simulations for the tumor cell density from the FK model (dotted lines), compared to the cell density predicted by the full model (solid line). Results from the FK model are obtained using different values for the tumor proliferation and diffusion rates (see Table S1).

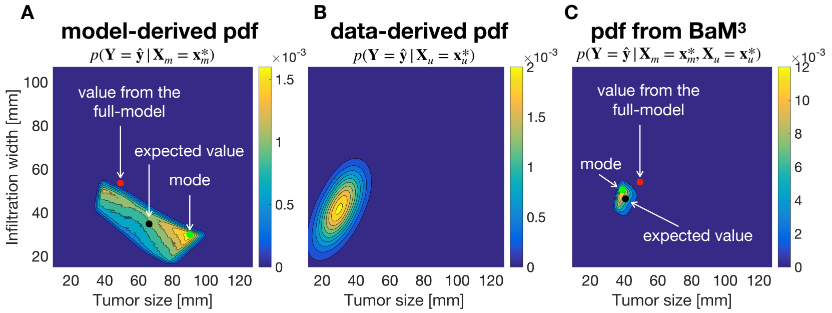


Figure S3: **A-C** Application of the BaM$^3$ method to a representative patient. The clinical presentation time $t_0$ is of 24 months and the prediction time $t_p$ is 12 months. **A** Pdf obtained from the FK model, plotted over the (TS, IW) space. The clinical output predicted by the full model (red dot), the expected value of the distribution (black dot), and its mode (green dot) are displayed before and after application of the BaM$^3$ method. **B** Data-driven pdf for the specific patient calculated through the kernel density estimator trained over the patient ensemble. **C** Probability distribution obtained from the BaM$^3$ method.

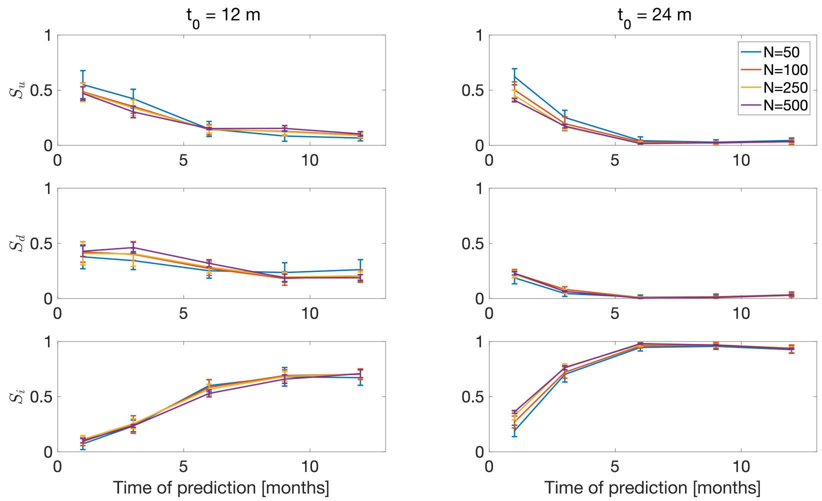


Figure S4: Prediction scores for different patient numbers $N$. The ratio of predictions that were unchanged ($S_u$), deteriorated ($S_d$) or improved ($S_i$) by the BaM$^3$ method are compared at different prediction times $t_d$ and different clinical presentation times $t_0$. The error bars represent the standard deviation of the results, obtained after 10 realizations of the same condition.

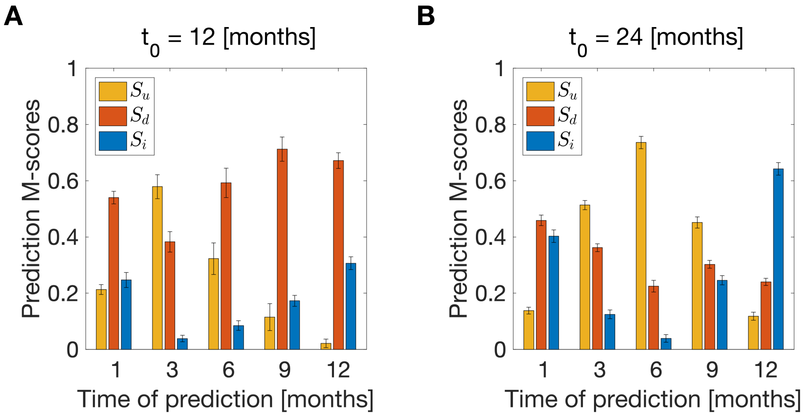


Figure S5: Prediction scores in terms of the ratio of predictions that have been unchanged ($S_u$), deteriorated ($S_d$) or improved ($S_i$) by the BaM$^3$ method. In these plots, we use the distribution mode to calculate the relative errors in the predictions for clinical presentation times $t_0$ of 12 and 24 months (**A**, **B**, respectively). The case shown refers to a number $N$ of patients of N=500. The error bars represent the standard deviation of the results, obtained after 10 realizations.

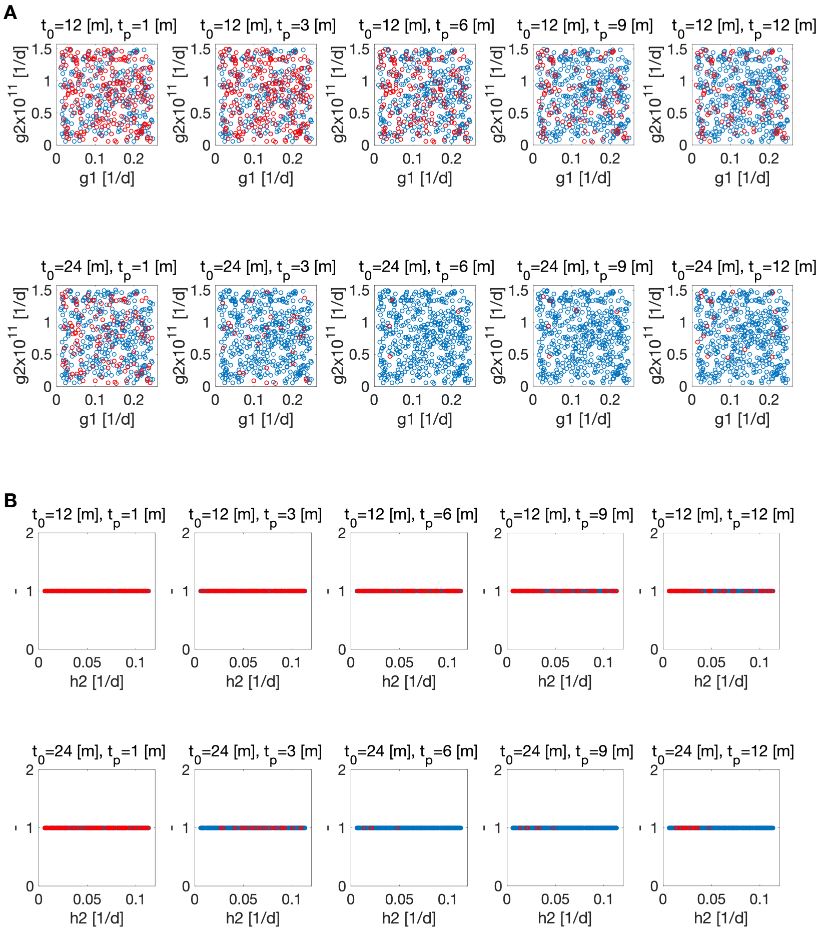


Figure S6: Scatter plots showing the distribution of patients for which the method fails to improve the model predictions (red dots). Each plot represents a different prediction time $t_p$ at a specific presentation time $t_0$. **A** Scatter plots for the parameter couple $g_1$ and $g_2$, controlling vascular formation and occlusion rate, respectively. **B** Scatter plots for the parameter $h_2$, which controls oxygen consumption by tumor cells.

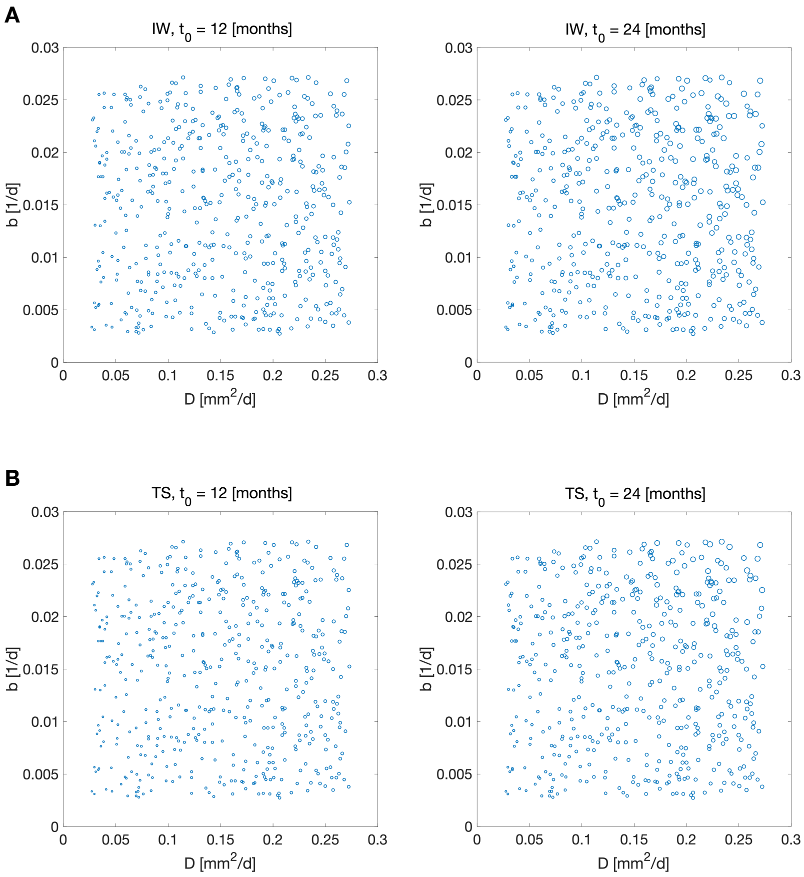


Figure S7: Scatter plots showing the distribution of IW and TS among different patients at the two clinical presentation times $t_0 = 12, 24$ months. The area of each dot is proportional to the corresponding value of IW or TS.

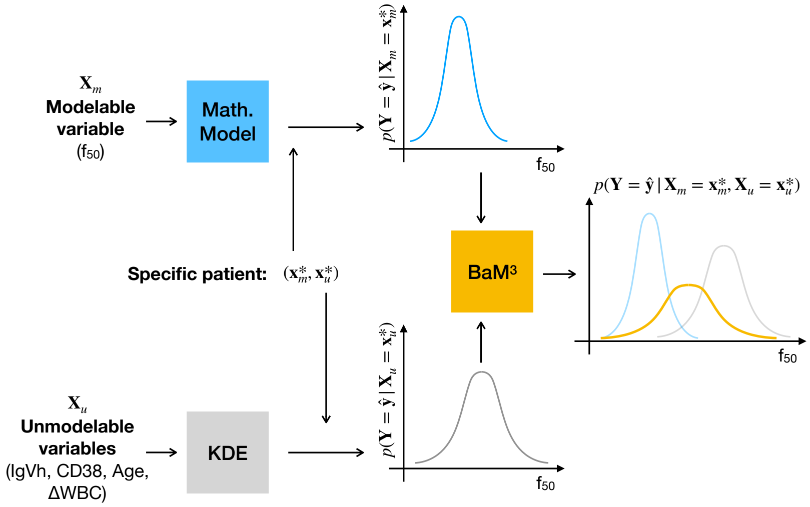


Figure S8: Schematic of approach used for the CLL patients. The mathematical model provides the pdf for the clinical outcome ($\mathbf{Y}$) having as input the modelable variable ($\mathbf{X}_m$). Note that in this case both the clinical output and the modelable variable correspond to the fraction of labeled cells at day 50 ($f_{50}$). At the same time, KDE provides the pdf for the clinical output using a set of unmodelable variables ($\mathbf{X}_u$). After entering the data for a specific patient ($\mathbf{x}_m^*, \mathbf{x}_u^*$), the BaM$^3$ method is applied.

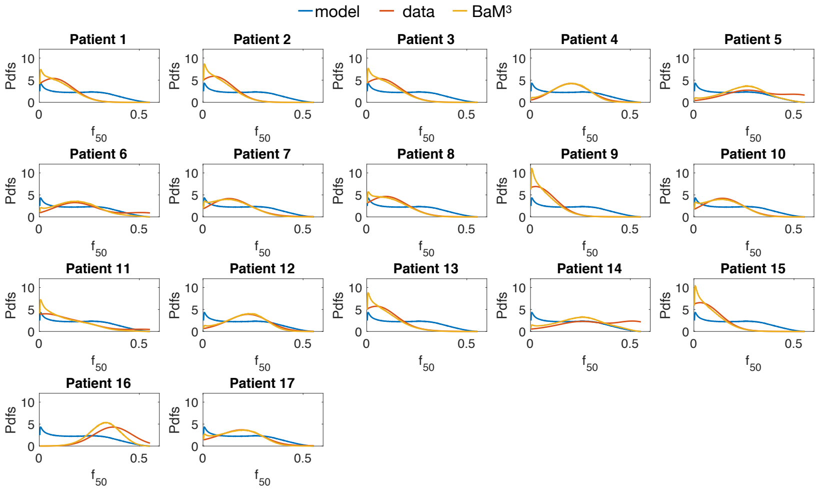


Figure S9: Probability distribution functions for the different patients in the CLL case. Here all the four unmodelables quantities have been used to inform the BaM$^3$ predictions.

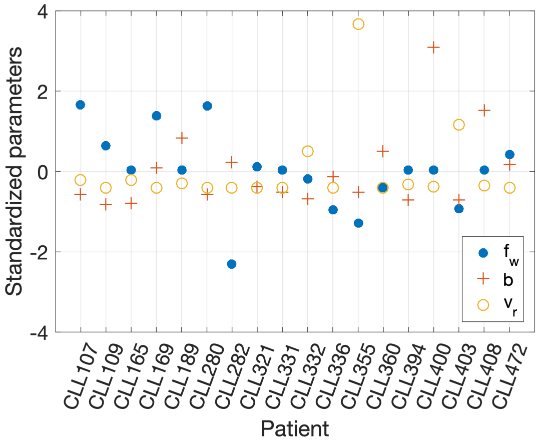


Figure S10: Distribution of the standardized parameters for the different patients in [6]. To standardize the parameters, we calculated the mean and standard deviation of each parameter group (i.e. $f_w, b, v_r$). Then, for each parameter value, we subtracted the corresponding mean and divided by the corresponding standard deviation.

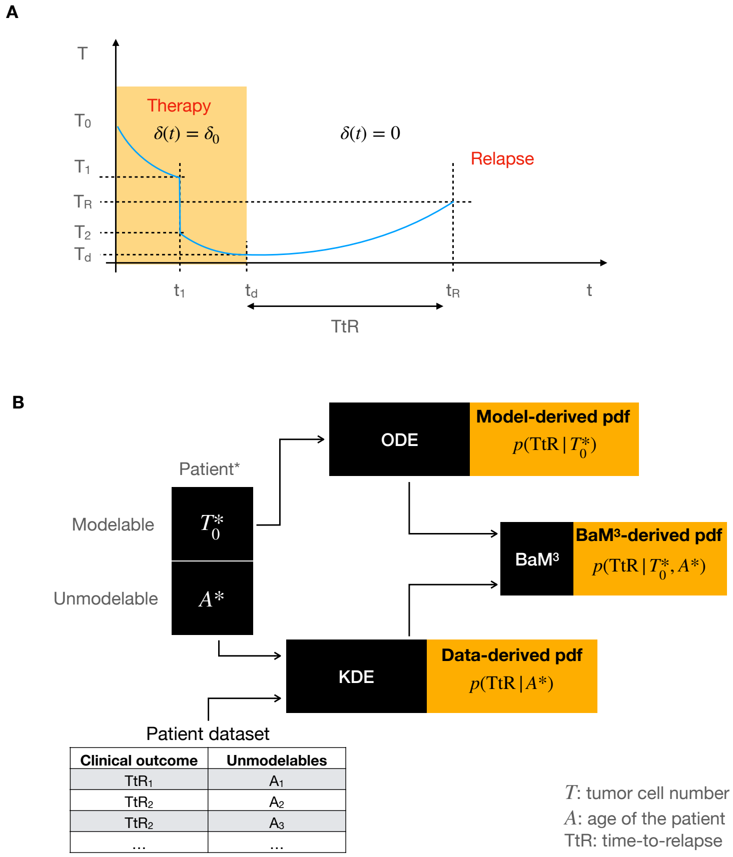


Figure S11: **A** Therapy schedule and tumor regrowth in the ovarian cancer case. The initial tumor cell number is denoted by $T_0$. Neoadjuvant chemotherapy is administered to patients between $0 < t < t_1$, then surgery is performed and the tumor reduces from $T_1$ to $T_2$. Then, another round of chemotherapy is administered, from $t_1$ to $t_d$. After therapy, growth resumes until the tumor reaches the relapse size $T_R$. The time that the tumor cell number takes to reach $T_R$ starting from $t_d$ is the clinical output of the problem, the time-to-relapse (TtR). **B** Schematic for the BaM$^3$ method applied to the ovarian cancer dataset. The modelable and unmodelable variables are given by $T$ and $A$, the tumor cell number and patient age at diagnosis, respectively. The patient specific tumor cell number $T^*$ is used in the mathematical model block (named ODE, in the Figure), to build the corresponding pdf for the TtR. In turn, the patient specific age at diagnosis $A^*$ is used in a density estimation procedure (KDE block, in the Figure) to derive the data-driven pdf. Both model- and data-derived pdf are then used in the BaM$^3$ method to provide the probability distribution of the clinical output, given the patient specific information on the modelable and unmodelable variables.

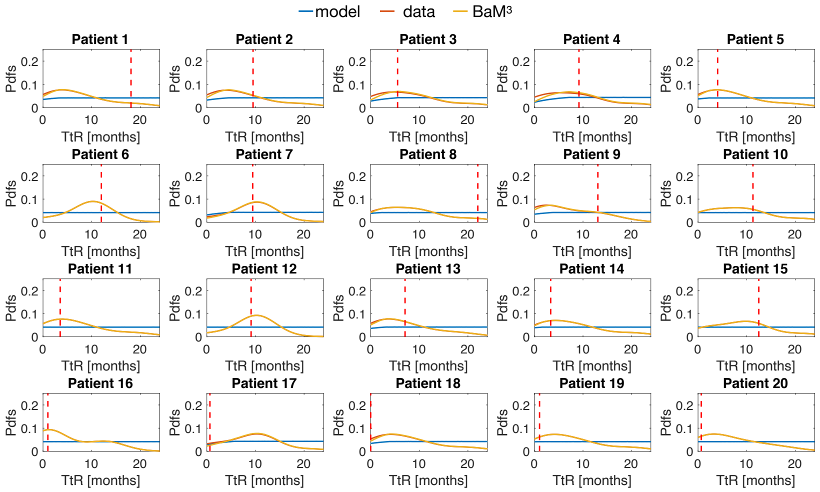


Figure S12: Probability distribution functions for the different patients in the ovarian cancer case, considering a uniform distribution of $\delta_0$ from 0.1 to 10 $d^{-1}$ in the mathematical model. This pdf refer to the 'uninformative' case.

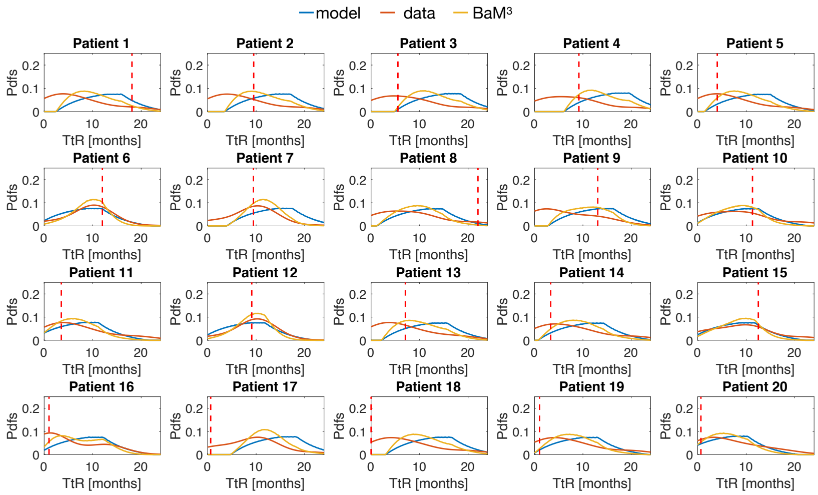


Figure S13: Probability distribution functions for the different patients in the ovarian cancer case, considering a uniform distribution of $\delta_0$ centered around the mean value obtained from the model fit (variation around the mean of $\pm 40\%$). This pdf refer to the 'fitted' case.

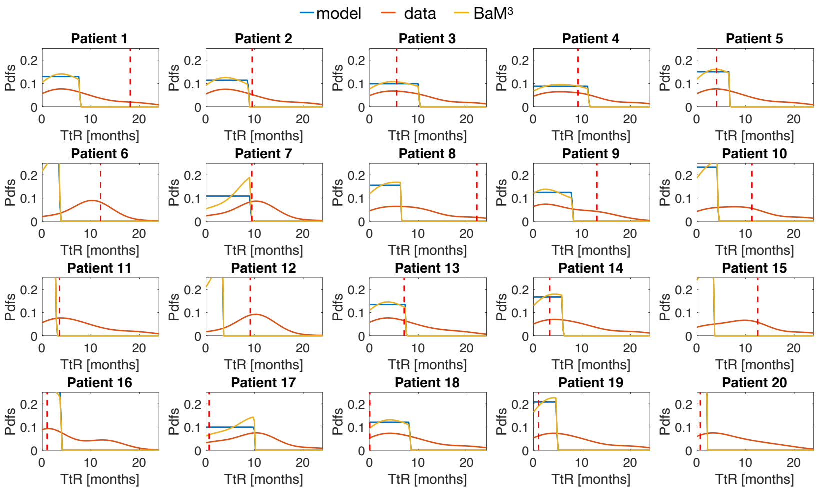


Figure S14: Probability distribution functions for the different patients in the ovarian cancer case, obtained by considering a simplified mathematical model.

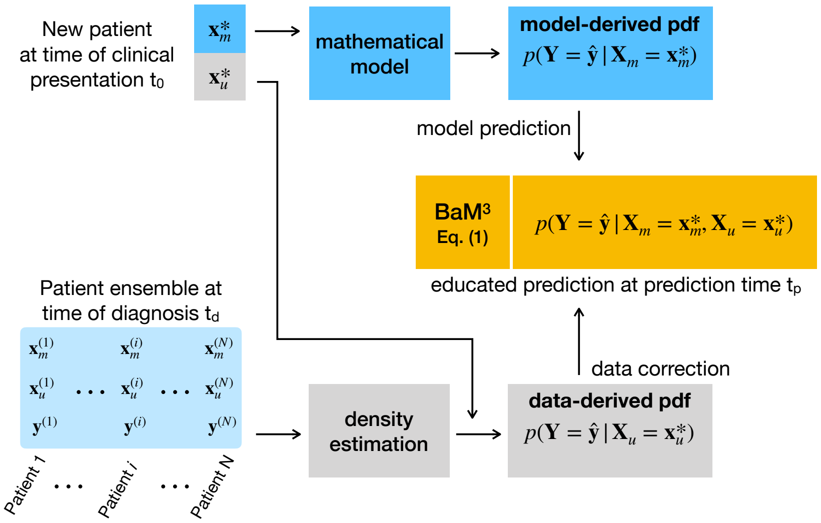


Figure S15: Schematic representation of the proposed methodology. The patient ensemble is constituted by modelable ($\mathbf{x}_m$), unmodelable variables ($\mathbf{x}_u$) and clinical observations $\mathbf{y}$ at the time of diagnosis $t_d$. Given a new patient ($\mathbf{x}_m^*$,$\mathbf{x}_u^*$) at the clinical presentation time $t_0$, the goal of the suggested approach is to find an estimate for the clinical observables $\hat{\mathbf{y}}$ at the prediction time $t_p$. The modelable variables $\mathbf{x}_m^*$ are used to setup the mathematical model, and provide predictions for the clinical observables $p(\mathbf{Y} = \hat{\mathbf{y}}|\mathbf{X}_m = \mathbf{x}_m^*)$. The unmodelable variables $\mathbf{x}_u^*$ are used in a density estimation method to provide the probability distribution function (pdf) $p(\mathbf{Y} = \hat{\mathbf{y}}|\mathbf{X}_u = \mathbf{x}_u^*)$, which is used to correct the predictions from the mathematical model. The proposed approach (Bayesian combination of mathematical modeling and machine learning, BaM$^3$) results in a new pdf of the clinical observables at time $t_p$, combining the outputs of the mathematical model and density estimation method.
